# Immune cell regulation in stem cell niche contacting glioblastomas

**DOI:** 10.1101/2022.02.16.480731

**Authors:** Todd Bartkowiak, Sierra M. Lima, Madeline J. Hayes, Akshitkumar M. Mistry, Justine Sinnaeve, Nalin Leelatian, Caroline E. Roe, Bret C. Mobley, Kyle D. Weaver, Reid C. Thompson, Lola B. Chambless, Rebecca A. Ihrie, Jonathan M. Irish

## Abstract

Glioblastomas (GBM) are tumors for which immune-targeted therapies have failed to show clinical benefit and for which few biomarkers provide context for meaningful therapeutic stratification. Radiographic contact of GBM tumors with the lateral ventricle stem cell niche correlates with worse patient prognosis; however, the extent to which proximity to the ventricle impacts antitumor immunity remains unknown. We demonstrate that T cell checkpoint receptor expression is elevated in ventricle-contacting GBM as is the abundance of a specific, suppressive CD32^+^CD44^+^HLAD^high^ myeloid population suggesting a distinct immunoregulatory influence on antitumor immunity in proximity to the lateral ventricle. Phospho-specific mass cytometric profiling revealed extensively impaired immune signaling in ventricle-contacting GBM in response to inflammatory cytokine stimulation, further supporting a suppressive milieu influencing immunity at the lateral ventricle. Collectively, we identify a regulatory impact of ventricle contact on antitumor immunity in the brain, and reveal novel clinically targetable mechanisms of immunomodulation in patients with glioblastoma.

**Significance Statement:** We demonstrate that the immune microenvironment of glioblastoma tumors contacting the lateral ventricle differs from non-contacting tumors. This work connects immune-biology to a radiographically detectable feature, the lateral ventricle, and highlights non-invasive imaging as a means to identify targetable immune features in glioblastoma tumors.

## Introduction

Glioblastoma (GBM) is the most common primary brain tumor, accounting for up to 60% of all brain tumors (1). Despite standard-of-care chemo/radiotherapy, median overall survival remains 15-months post diagnosis (2). While efforts have been made to phenotypically and molecularly characterize GBM tumors and illuminate the surrounding stromal elements impacting gliomagenesis, few cell-intrinsic factors beyond isocitrate dehydrogenase (*IDH*) mutation and O-6-methylguanine-DNA methyltransferase (*MGMT*) promoter methylation have proven effective at stratifying patient outcome and guiding clinical care (3). Additional, less-invasive, risk-stratifying features are urgently needed to make headway in understanding the complex cellular milieu in the tumor and advance therapeutics in the clinic.

The majority of GBM tumors present in the cerebrum, and although GBM may arise anywhere within the brain parenchyma, both adult and pediatric patients with primary high-grade gliomas (HGG) that exhibit radiographic contact with the lateral ventricles (LV) have worse prognosis (4, 5). This effect is independent of other predictive factors, such as patient age, performance status, or molecular characterization (6). While regional tumor position stratifies prognosis, what factors proximal to the LV contribute to poor prognoses remains unclear.

Immune cells comprise a major component of the tumor lesion and play a critical role in controlling tumor growth in both solid and hematologic malignancies. The immune microenvironment within the brain, however, is distinct from peripheral tissue and the antitumor potential of leukocytes in this discrete environment remains poorly characterized (7). Cytometric profiling has demonstrated regional differences in resident immune cell phenotypes under homeostatic conditions, particularly in the LV (8); however, how regional neuro-immunity is impacted in the context of primary tumor lesions in the ventricle remains poorly understood.

In this study, we performed multi-dimensional single-cell mass cytometry on 25 primary human GBM tumors demonstrating radiographic contact with the lateral ventricle (C-GBM) or located distally from the ventricle (NC-GBM) to comprehensively profile the immune infiltration in tumors within each region. Supervised and unsupervised machine learning approaches including clustering tools FlowSOM (9) and Root-mean-square deviation (RMSD) (10), population identification via Marker Enrichment Modeling (MEM) (11), and patient stratification using Citrus(12) and Risk Assessment Population Identification (RAPID) (13) identified distinct immune subsets enriched in C-GBM and NC-GBM that correlated with patient outcome. Moreover, several targetable immune receptors were elevated in C-GBM tumors suggesting that an immunosuppressive environment lies proximal to the LV. Lymphocytes and tissue-resident microglia were enriched in NC-GBM tumors and correlated with more favorable outcome, whereas anti-inflammatory M2-like monocyte-derived macrophages (MDMs) bearing a distinct CD44^+^CD32^+^HLA-DR^+^ phenotype and exhausted PD-1^+^TIGIT^+^ T cells were enriched in C-GBM tumors.

Phospho-specific mass cytometry revealed broad T cell signaling defects and differential use of myeloid signaling networks employed by C-GBM and NC-GBM leukocytes in response to inflammatory cytokine stimulation. CD32^+^HLA-DR^+^ MDMs were highly responsive to cytokines compared to their CD32^-^HLA-DR^-^ counterparts. Further, STAT3 phosphorylation dominated signaling networks in C-GBM immune infiltrates compared to NC-GBM infiltrates, highlighting a potential STAT3 driven immunosuppressive mechanism in C-GBM tumors. This work highlights differing immune microenvironments within MRI-defined regional tumor classes, suggests distinct, targetable mechanisms of immune dysregulation in GBM tumors in relation to the lateral ventricle, and emphasizes the potential for radiographic image-guided patient stratification methods to inform clinical care.

## Results

### 1. Peripheral immune cells are abundant in lateral ventricle contacting and non-contacting glioblastomas

We utilized high-dimensional mass cytometry to compare the immune composition of 25 freshly resected glioblastoma tissues either contacting the lateral ventricle (C-GBM; n=13) or distant from the ventricle (NC-GBM; n=12) (**Figure 1a**). All patients expressed the wildtype *IDH1/2* variant. The median overall survival was 366 days (range: 57-1588 days), typical of the disease. A full description of patient characteristics is given in **Supplementary Table 1**. As in prior reports (4), ventricle contact was a predictive factor of outcome. Resected tumor tissue was dissociated into single-cell suspensions and profiled using a 33-marker panel to characterize 13 expert-identified infiltrating immune subsets (**Figure 1b, Supplementary Table 2**).

**Figure 1:**
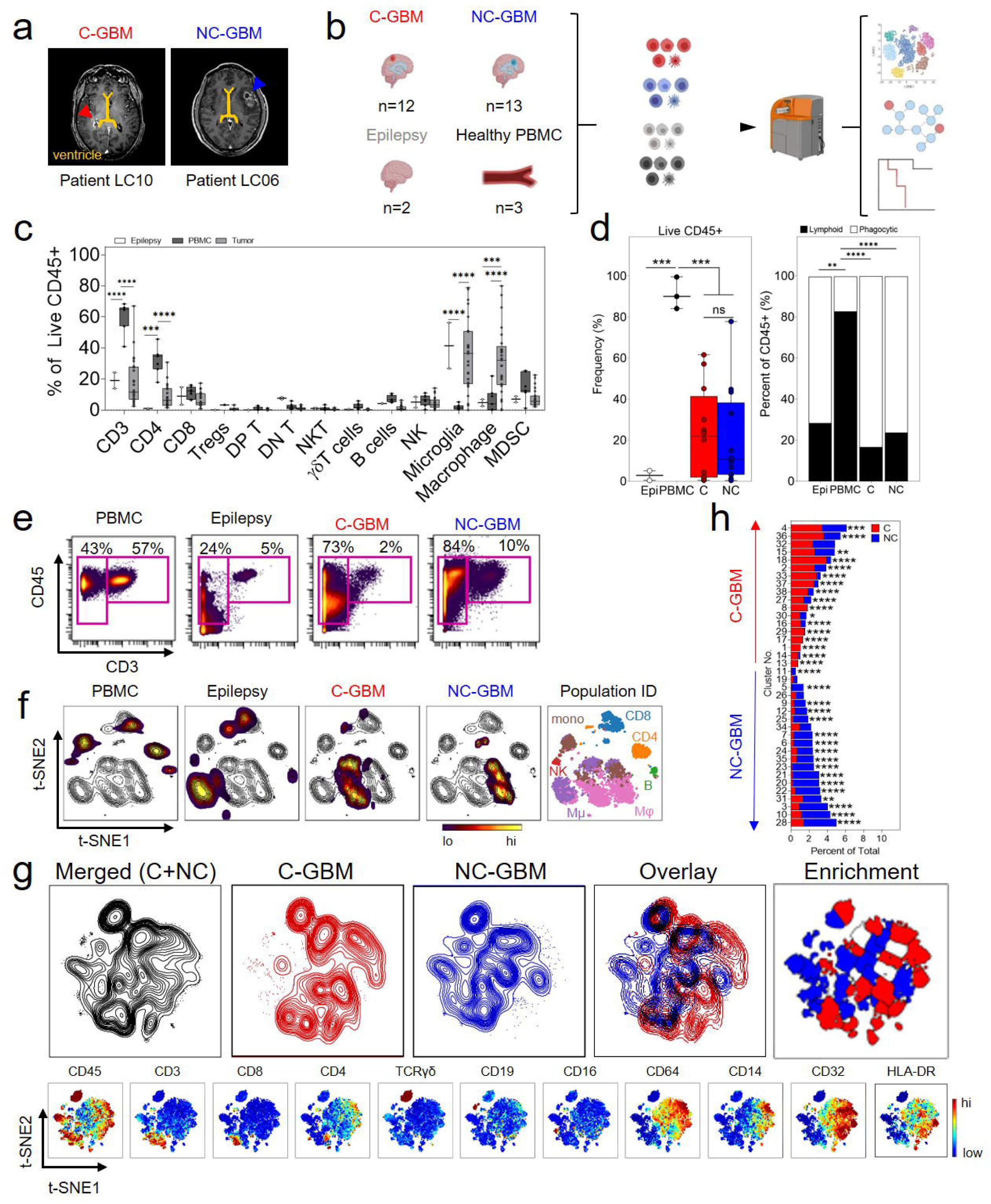
Lateral ventricle contacting and non-contacting glioblastomas are enriched in distinct immune subsets. **a)** Representative MRI radiographs of WHO grade IV glioblastoma tumors with confirmed contact with either of the lateral ventricles (left, C-GBM) or lacking ventricular involvement (right, NC-GBM). Yellow line indicates the lateral ventricle and arrows indicate the tumor mass. **b)** Tissue was obtained from patients with radiographically confirmed ventricle contacting (C-GBM, red) (n=12) or non-contacting (NC-GBM, blue) (n=13) glioblastomas, temporal lobectomy of parenchymal tissue from epileptic patients (light grey) (n=2) or peripheral blood mononuclear cells (PBMC, dark grey) from healthy volunteers. Single-cell suspensions were processed, and mass cytometry was performed before multiple high-dimensional analysis platforms assessed the immune phenotypes. **c)** Bar graphs demonstrating the frequency of expert-gated immune populations in epileptic brain (white), healthy donor PBMC (dark grey), or glioblastoma tumors (light grey). **d)** Bar graphs demonstrating the total leukocyte infiltration (left) and lymphoid and phagocyte fractions (right) in epileptic brain (Epi), healthy donor blood (PBMC), C-GBM (C) or NC-GBM (NC). **e)** Representative biaxial plots demonstrating the frequency of CD45^+^CD3^+^ T cells and CD45^+^CD3^-^ non-T cells in PBMC, epileptic brain tissue, C-GBM and NC-GBM tumors respectively. **f)** Representative t-SNE plots of all CD45^+^ leukocytes identified in the blood, non-tumor bearing brain, or GBM tumors (heat) overlaid onto all CD45^+^ events. Multi-color overlay represents expert gated populations overlaid onto the t-SNE axes. **g)** Live CD45^+^ cells were merged from all GBM patients (black contour), patients with C-GBM tumors (red contour), or NC-GBM tumors (blue contour). Overlaid t-SNE plots indicate areas of immune infiltration unique to LV contacting or non-contacting patients. Enrichment indicates which computationally gated immune populations were statistically enriched in LV-contacting or non-contacting gliomas. Heat displayed for various markers indicates phenotypes for major immune subsets. **h)** Bar graphs demonstrating the frequency of immune cells found within each computational cluster as a percent of total CD45+ leukocytes. Bars indicate median ± IQR. * = p<0.05, ** = p<0.01, *** = p<0.001, **** = p <0.0001.

We first compared the tumor immune composition to that of healthy donor peripheral blood mononuclear cells (PBMCs) or resected epileptic brain tissue (**Figure 1c**). Consistent with prior reports, T cell infiltrate in GBM, particularly the CD4 T cells, was similar to epileptic brain, accounting for 11% and 6% of the total leukocyte infiltrate respectively. Other lymphocyte populations—Tregs, CD4^+^CD8^+^ (DPT), CD4^-^CD8^-^ (DNT), natural killer (NK) T cells, γδ T cells, B cells and NK cells—each made up <5% of the leukocyte fraction in the tumor. CD45^low^CD64^+^ microglia were the most abundant leukocyte population in the brain accounting for 36% of the immune fraction in GBM tumors. CD45^high^CD14^+^ peripheral monocyte-derived macrophages (MDMs) were the next most abundant leukocyte population, accounting for 32% of leukocytes in glioma, compared to 4% in PBMC and 5% in epileptic brain, consistent with an extensive peripheral myeloid infiltrate in glioblastomas (14).

Considering that the extent of lymphocytic infiltration correlates with more favorable outcomes in peripheral solid tumors (15), we hypothesized that total leukocyte abundance in NC-GBM may drive favorable outcomes observed in this cohort. While the total leukocyte abundance in C-GBM and NC-GBM was higher than epileptic brain, no significant difference was found between the two patient cohorts, accounting for 23% (range: 0.47-61.62%) and 20% (range: 0.38-77.87%) of the tumor mass respectively (**Figure 1d**). Moreover, the ratio of lymphoid cells (T, B, NK) and phagocytes (MDM and microglia) was not significantly different between C-GBM and NC-GBM, suggesting that neither total leukocyte abundance nor bulk lymphocyte abundance fully accounts for differences in survival outcome between C-GBM and NC-GBM patients. However, meaningful differences in the frequency and phenotypes of specific lymphocyte and phagocyte populations were evident in C-GBM and NC-GBM tumors. For instance, the abundance of CD3^+^ T cells was elevated in NC-GBM compared to C-GBM or epileptic brain tissue (**Figure 1e**). By overlaying leukocytes onto t-SNE axes, we identified distinct leukocyte populations in PBMCs, epileptic brain, and GBM tissue. The phenotypes of macrophages in C-GBM and NC-GBM tumors differed, with macrophage populations occupying distinct islands within the t-SNE space (**Figure 1f**), suggesting that phenotypic differences in immune infiltration correlated with ventricle contact status.

To further explore the extent of these cohort-level differences in immune abundance, data from 19 GBM patients (9 C-GBM, 10 NC-GBM) was used to investigate the immune composition of C-GBM and NC-GBM tumors. Live CD45^+^ cells were equally sampled from each patient prior to plotting on common t-SNE axes using 33 measured dimensions (**Figure 1g**). Differences in immune cell composition were evident when comparing the distribution of C-GBM and NC-GBM samples across the common axes. FlowSOM clustering on the t-SNE axes defined 38 phenotypically distinct immune cell clusters across the entire patient cohort (**Figure 1g, Supplementary Figure 1a**). The majority of patients were well represented, with between 2 and 17 patients contributing to each cluster (**Supplementary Figure 1b).** Of the 38 clusters, 17 were statistically enriched in immune cells from C-GBM patients, and 17 were enriched in cells from NC-GBM patients. Four clusters (19, 26, 32, 34) were not statistically enriched in either cohort (**Figure 1g, Figure 1h, Supplementary Figure 1**). Taken together, these data highlight extensive differences in the immune composition of GBM tumors and suggest that while immune infiltration occurs in both C-GBM and NC-GBM, the relative composition of the immune cell fraction in ventricle contacting tumors is distinct from that of ventricle non-contacting tumors.

### 2. Five immune cell subsets are differentially enriched in ventricle contacting and non-contacting glioblastomas

We next used machine learning to perform unbiased computational analysis of the immune microenvironment of GBM tumors and identify the most distinct immune subsets associated with C-GBM and NC-GBM tumors. Using the Citrus algorithm to compare leukocyte phenotypes and frequencies between our two cohorts (12), we identified 10 immune clusters differentially enriched in C-GBM and NC-GBM tumors. Five terminal clusters representing the most phenotypically distinct populations were selected for further study (**Supplementary Figure 2a**). Three Citrus clusters were enriched in NC-GBM (Clusters 1, 2, and 5), while 2 clusters (Clusters 3 and 4) were enriched in C-GBM (**Figure 2a**). The identities of these clusters inferred from Citrus were confirmed via traditional biaxial gating and computational labeling using marker enrichment modeling (MEM) (11) (**Figure 2b-f**). Cluster 1, enriched in NC tumors, consisted of γδ T cells and CD3^+^ CD4^-^ CD8^-^ T cells (DNT) previously identified in GBM (16). Cluster 2 was a population of resident microglial cells characterized as CD45^low^CD64^+^CD14^-^HLA-DR^+^CD32^+^ (**Figure 2c**). Cluster 5, also enriched in NC tumors, consisted of lymphocyte populations: CD8 T cells (28%), B cells (21%), NK cells (11%), and CD4 T cells (7%) (**Figure 2f**). In contrast, Clusters 3 and 4, which were enriched in C-GBM, were characterized as CD45^high^CD64^+^CD14^+^, consistent with peripheral monocyte-derived macrophages (MDMs) (17) (**Figure 2d, e**). Expert-guided gating on patient samples not included in the initial analysis (n=6) identified similar immune phenotypes consistent with the Citrus findings and MEM labels (**Supplementary Figure 2b-d**), and confirmed statistical enrichment of the immune populations identified by the Citrus algorithm (**Supplementary Figure 2e**).

**Figure 2:**
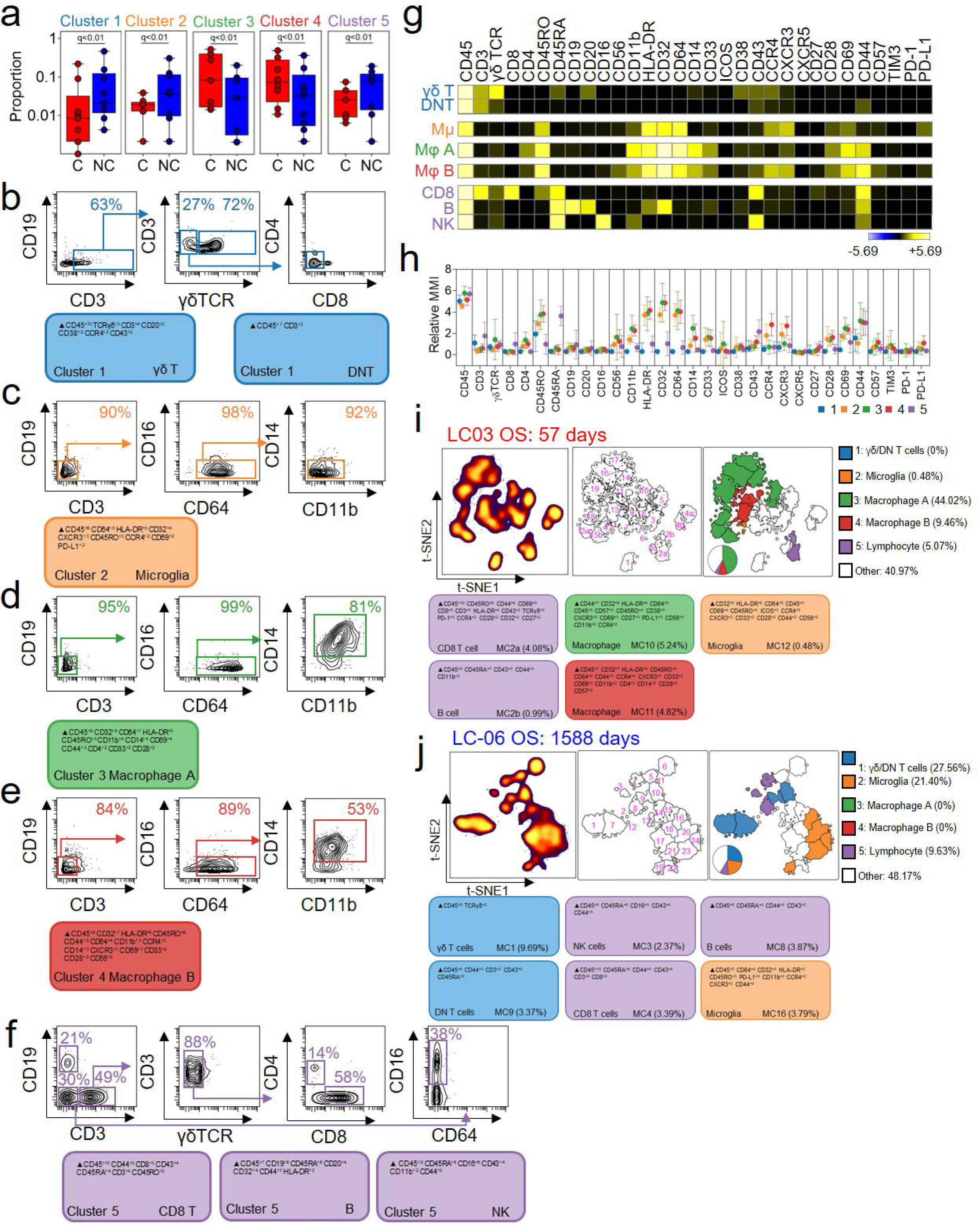
Differential enrichment of five immune phenotypes distinguish ventricle contacting and non-contacting glioblastoma. **a)** Citrus clustering of live CD45+ leukocytes in the tumor microenvironment of C-GBM and NC-GBM tumors revealed differential enrichment of five immune subsets. Representative biaxial gating of pooled Citrus clusters and Marker Enrichment Modeling (MEM) population identification classified each cluster as **(b)** γδ and CD4^-^CD8^-^ DNT cells, **(c)** microglia, **(d)** and **(e)** macrophages and **(f)** lymphocyte populations of CD8 T cells, B cells and NK cells. **(g)** Heatmaps of each phenotypic marker used to classify each immune subset reveal the expression levels of each immune receptor. **(h)** Quantification of the arcsinh transformed expression level of each immune marker within each subset. Representative t-SNE plot of all CD45+ leukocytes infiltrating a C-GBM tumor **(i)** or NC-GBM tumor **(j)**. Cell density (left t-SNE), FlowSOM clustering on the t-SNE axes (middle), and Citrus overlay and quantification (right) determined the relative frequency of each immune cell subset within each patient sample. A subset of color-coded MEM labels identified the phenotype and frequency of key Citrus-identified immune populations. In **(a)**, a regularized regression model was used as a final step in the Citrus analysis to identify stratifying clusters. Predictive analysis of microarrays (PAM) stratified immune clusters. A False Discovery Rate <1% (q) was used to determine significance in all instances.

Examining each cluster more deeply, γδ T cells in Cluster 1 expressed low levels of CD45RA, CD38, CD43, and CCR4, while DNT cells expressed CD43 and CD44, consistent with an effector T cell phenotype (**Figure 2g, Figure 2h**). Similarly, CD8 T cells present in Cluster 5 expressed high levels of CD43, CD44, and CD45RA indicative of activated CD45RA expressing effector memory T cells (T_EMRA_) (18). B cells in NC-GBM tumors expressed low levels of HLA-DR and high levels of the inhibitory Fc receptor CD32 (FcγRII), suggesting impaired antigen presentation capacity. NK cells within Cluster 5 were characterized as CD16^+^CD56^-^CD43^+^CD44^+^CD11b^low^, a mature, cytotoxic NK cell phenotype. Finally, microglia enriched in NC tumors (Cluster 2) were characterized as CD45RO^low^CD11b^low^HLA-DR^+^CD32^+^CD64^+^CCR4^low^CD69^low^PD-L1^low^, characteristic of activated microglia. In contrast, MDMs within contacting tumors (Clusters 3 and 4) expressed higher levels of CD45RO, CD11b, CD32, and CD44 than their microglial counterparts. The distinguishing characteristics of macrophage populations A (Cluster 3) and B (Cluster 4) included higher expression of CD14 in Cluster 3 and higher expression of the chemokine receptors CXCR3 and CCR4 in Cluster 4 (**Figure 2g, h**), confirming that these populations represent two activated, phenotypically distinct myeloid subsets in the C-GBM microenvironment.

Citrus events were then overlaid on patient-specific t-SNE maps (**Supplementary Figure 3**) to determine the source of subsampled Citrus clusters and compare the phenotypes and frequencies of immune cells within larger FlowSOM subsets (13). Immune phenotypes identified in the subsampled set of cells were no statistically different than the larger immune populations (p=0.6844) (**Supplementary Figure 3a-d**). Consistent with the abundance of total CD45^+^ populations discussed above, the average number of phenotypically distinct FlowSOM clusters, an estimate of overall immunological diversity, was similar between C-GBM and NC-GBM tumors (23 vs. 22 FlowSOM clusters respectively), as was the number and proportion of clusters represented by Citrus in the entire dataset (**Supplementary Figure 3e**), suggesting that the degree of diversity within the immune infiltrate does not significantly contribute to outcomes associated with ventricle contact status.

The frequency and immune phenotype of individual patients’ clusters defined by MEM labels (**Figure 2i, Figure 2j, Supplementary Figure 3f, Supplementary Figure 4a-b**) reflected the larger patterns identified by Citrus. For example, Patient LC03 who had a C-GBM tumor and the worst overall survival (57 days post resection), possessed abundant frequencies of macrophages corresponding to Citrus cluster 3 (44%; 8/23 clusters) and cluster 4 phenotypes (9.46%; 2/23 clusters). Five percent of the immune fraction in this patient consisted of CD8 T cells (4%) and B cells (1%) with <1% of the microglia and γδ/DNT populations (Clusters 1 and 2) (**Figure 2i, Supplementary Figure 4a**). Conversely, Patient LC06, who had an NC-GBM tumor and the greatest overall survival (1588 days post resection) included no macrophages in clusters 3 or 4, but had an abundance of microglia (21%), DNT cells (17.87%), γδ T cells (9.69%), NK cells (2.37%), CD8 T cells (3.39%), and B cells (3.87%), collectively reflecting phenotypes found in clusters 1,2, and 5 (**Figure 2j, Supplementary Figure 4b**). We next sought to compare the phenotypes of all FlowSOM immune clusters in order to identify common immune populations across all patients in our cohort. Comparison of all clusters identified by FlowSOM (455 clusters from all patients) using Root-mean-square-deviation (RMSD) identified populations enriched in C-GBM or NC-GBM tumors overlapping with Citrus phenotypes (**Supplementary Figure 5a**). Across all patients within the cohort, RMSD analysis identified 17 common immune phenotypes across our samples. The majority of RMSD clusters were well represented across the cohort, with RMSD cluster 6 (CD8 T cells) the most represented cluster, and RMSD cluster 12 (microglia) the least represented (**Supplementary Figure 5b**). The abundance of each cluster within each patient sample (**Supplementary Figure 5c**), the statistical enrichment (**Supplementary Figure 5d**), and MEM labels for each immune phenotype (**Supplementary Figure 5e**) were consistent with Citrus findings.

Taken together, these data indicate that although individual patients possessed a range of immune cell phenotypes in their tumors, common immune signatures could be identified across the cohort. In particular, microglia and lymphocytes, specifically γδ T cells, DNT cells, CD8 T cells, NK cells, and B cells were enriched in patients with NC-GBM. Conversely, patients with C-GBM possessed an immune microenvironment enriched in MDMs, revealing starkly contrasting immune microenvironments in C-GBM and NC-GBM tumors.

### 3. Immune cell frequencies correlate with patient outcome

We next determined if the relative abundance of immune populations correlated with outcome, similar to observations in peripheral solid tumors (19). Using a Cox proportional hazards model, γδ T cells and DNT cells (Citrus cluster 1) and macrophages (Citrus clusters 3 and 4) did not significantly correlate with patient outcome (**Figure 3a**). Microglial cells (Citrus cluster 2) and lymphocytes (Citrus cluster 5), however, correlated with improved patient survival. Higher frequencies of microglia cluster 2 (>2%) correlated with more favorable overall survival outcomes (median 560.5d vs 252.0d; p=0.0405, HR= 0.3782, CI [0.1414-1.012]) as did lymphocytes (>6%) (median 507d vs 215d; p=0.0126, HR=0.3226, CI [0.155-0.9015]). Importantly, 5/6 patients in the “microglia high” and 6/7 patients in the “lymphocyte high” group presented with NC tumors, consistent with our previous findings (**Figure 2**).

**Figure 3:**
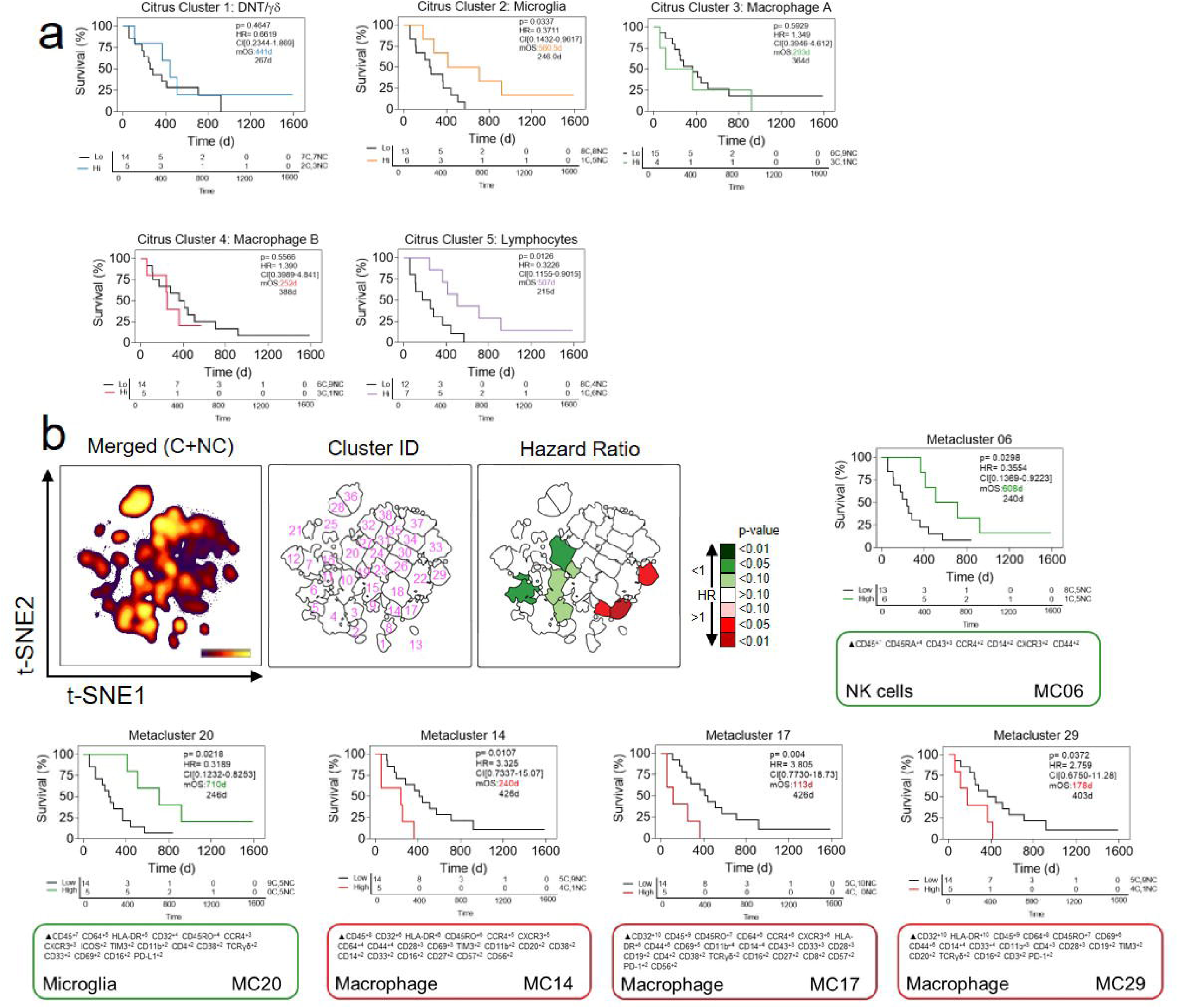
Immune subset frequencies correlate with patient outcome. **a)** Kaplan-Meier curves indicating overall survival in GBM patients with high vs low frequencies of Citrus-identified immune populations. **b)** Kaplan-Meier curves for overall survival in immune subsets stratifying patient outcome identified by RAPID analysis. T-SNE plots in b indicate the cell density (left), cluster number (middle) and p-value of the hazard ratio associated with the frequency of each cluster in the entire cohort. Calculated MEM labels identified key features of stratifying immune subsets.

To corroborate these results using an orthogonal, unsupervised computational approach, the Risk Assessment Population IDentification (RAPID) workflow was used to directly reveal immune cell clusters whose abundance correlated with patient outcome (13). RAPID identified eight populations that were either strongly (p<0.05) or modestly (p<0.1) correlated with patient outcome (**Figure 3b**). These phenotypes were stable through iterative runs of RAPID, including repeated subsampling (**Supplementary Figure 6**). Consistent with Citrus results, one statistically significant RAPID metacluster (06) represented a population of CD43^+^ NK cells whose abundance was predictive of favorable outcomes (p=0.0298, HR=0.3554, CI [0.1369-0.9223]). Five out of six patients with high frequencies of these NK cells presented with NC tumors, consistent with Citrus results. Greater frequencies (>1.32%) of RAPID metacluster 20 (CD45^int/low^CD64^+^HLA-DR^+^CD45RO^low^ microglia) also correlated with longer survival (710d vs 246d; p=0.0218, HR=0.3189, CI [0.1232-0.8253]). All five patients with high abundance of this microglial subset presented with NC-GBM tumors. Conversely, three subsets of CD45^high^CD64^+^CD32^+^HLA-DR^+^CD14^+^ macrophages correlated with poor prognosis (RAPID metaclusters 14, 17, and 29). For example, Metacluster 17, an MDM population phenotypically similar to Citrus Cluster 4, correlated with roughly 4-fold worse prognosis (median overall survival (mOS) = 113d vs 426d, p=0.004, HR= 3.805, CI [0.7730-18.73]). Critically, patients with the highest frequencies of MDMs presented with ventricle-contacting tumors, providing further evidence that blood-derived macrophages correlate with ventricle tumor contact and worse prognosis.

Collectively, these data suggest that infiltration by specific immune cells correlates with survival in glioblastoma and LV contact status. Orthogonal approaches confirmed that computationally identified subsets of microglial cells and lymphocytes correlated with more favorable outcomes whereas increased infiltration of subsets of peripheral macrophages correlated with worse outcome independent of LV contact status. Thus, C-GBM and NC-GBM have distinct immune microenvironments and contrasting patient outcomes that align closely with the expected functional role of the immune cells present in each.

### 4. The immune microenvironment of lateral ventricle contacting gliomas is enriched in inhibitory checkpoint receptor expression

We next sought to identify targetable immune receptors that may give insight into mechanisms of immunoregulation in tumors proximal to the ventricle and inform clinical decision making. We used machine learning to identify individual cellular features that were enriched in C-GBM or NC-GBM immune infiltrates (**Figure 4a**, **Supplementary Figure 7a**). Citrus identified elevated expression of 5 markers on 7 immune subsets in C-GBM tumors (**Figure 4a**, **Supplementary Figure 7a-b**). Of note, the checkpoint receptor PD-1 was elevated on both CD4 and CD8 T cells infiltrating C-GBM tumors, as were CD32, CD69, CD44, and HLA-DR on C-GBM infiltrating peripheral MDMs. Moreover, CD32 and CD69 were elevated across multiple phagocytic populations infiltrating C-GBM including MDMs and microglia. Lastly, HLA-DR was elevated on a population of γδ T cells infiltrating C-GBM tumors (**Figure 4b-c**). Consistent with Citrus results here and in Figure 2, expert-guided biaxial gating confirmed an increased frequency of CD32^+^CD44^+^HLA-DR^+^ macrophages infiltrating C-GBM (**Figure 4d**) compared to NC-GBM.

**Figure 4:**
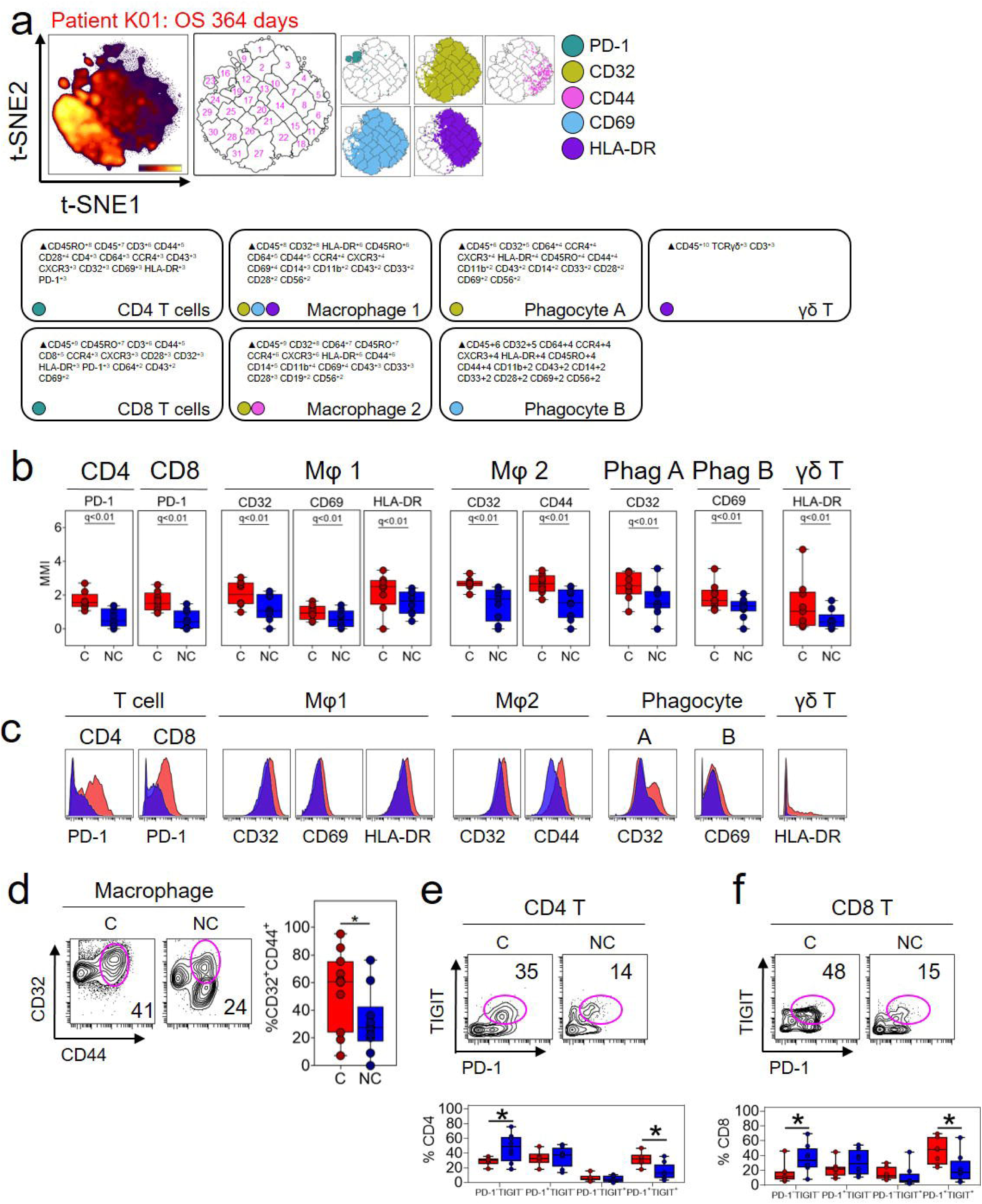
Immunosuppressive checkpoint receptors are enriched in ventricle contacting glioblastomas. **a)** Representative t-SNE plot indicating the density of all CD45+ leukocytes (left), FlowSOM clusters on the t-SNE axes, and overlaid immune populations with enriched expression of indicated immune markers. MEM labels indicate the cellular phenotype in which the indicated markers were differentially expressed. **b)** Box and whisker plots indicating the arcsinh transformed median expression values of indicated immune receptors within Citrus-identified populations of interest. **c)** Histograms of pooled patient Citrus clusters from C-GBM (red) and NC-GBM patients (blue). **d)** Representative biaxial plot (left) and box and whisker plots indicating the frequency of CD32^+^CD44^+^ macrophages identified by Citrus. **e**) Representative biaxial plots and box and whisker diagrams demonstrating the frequency of TIGIT and PD-1 co-expression in CD4 T cells **(e)** and CD8 T cells **(f)** infiltrating glioblastoma tumors. In **(a)**, a regularized regression model was used to as a final step in the Citrus analysis to identify stratifying clusters. Predictive analysis of microarrays (PAM) stratified immune clusters. A False Discovery Rate of 1% (q) was used to determine significance in all instances. Bars indicate median ± IQR. * = p<0.05, ** = p<0.01, *** = p<0.001, **** = p <0.0001.

Expert gating strategies confirmed increased frequencies of PD-1^+^ CD4, CD8, and DNT cells within C-GBM tumors (**Supplementary Figure 7c-l**, **Supplementary Tables 2 and 3**). Critically, PD-1^+^ T cells infiltrating C-GBM co-expressed the inhibitory receptor TIGIT at a higher frequency in C-GBM (46% vs. 23%), whereas a higher frequency of PD-1^-^ TIGIT^-^ T cells were found in NC-GBM (36% vs 17%), suggesting that C-GBM T cells may be more phenotypically exhausted compared to their NC-GBM counterparts (**Figure 4e, f**). Additionally, C-GBM tumors bore increased frequencies of T and NK cells (**Supplementary Figure 7c-f**), myeloid subsets (**Supplementary Figure 7g-i**) and B cells (**Supplementary Figure 7j**) with an activated phenotype compared to NC-GBM tumors. While CD45^-^ tumor stromal cells in C-GBM and NC-GBM expressed a variety of immune checkpoint receptors/ligands, we found no difference in the frequency of checkpoint positive tumor cells in C-GBM and NC-GBM tumors (**Supplementary Figure 7k**). Interestingly, increased frequencies of T cells expressing several markers (CD27, CD32, CXCR3, CCR7) suggested that these cells in C-GBM tumors possessed a phenotype consistent with immunological memory. Indeed, the frequency of CD45RO^+^CCR7^+^ central memory T cells (T_cm_) was elevated in C-GBM tumors (**Supplementary Figure 7l**). Further, consistent with Citrus results (**Figure 2**), increased frequencies of CD45RO^-^CCR7^-^ T_EMRA_ CD8 T cells were found in NC-GBM tumors.

Taken together, our data suggests that not only does tumor proximity to the lateral ventricle influence the types of immune cells infiltrating GBM tumors, but also impacts the phenotype, and potential function of these cells. Through multiplex machine learning and expert-guided strategies, we have identified multiple therapeutically targetable checkpoint receptors on distinct immune subsets in GBM tumors.

### 5. Immune receptor expression correlates with patient outcome

We next sought to identify which of the differentially enriched immune markers identified by Citrus may help stratify patient outcomes. To do so, we developed the median marker implementation of RAPID (mmRAPID) whereby the median intensity values for select markers in RAPID clusters were correlated with patient outcome (See Methods). Median expression values of 12 immune receptors across 46 immune clusters correlated with patient outcome (**Figure 5a**). Consistent with enriched immune receptor expression in C-GBM tumors (**Figure 4**) and immune abundance correlating with outcome (**Supplementary Figure 6**), a 2-fold increase in PD-1 expression in CD4 T cells (mmRAPID Cluster 9) (**Figure 5b**) and CD8 T cells (mmRAPID Cluster 25) (**Figure 5c**) correlated with 2-fold worse survival (215d vs. 441d and 240d vs 570d, respectively). Importantly, although contact status was not used in the mmRAPID analysis, 6/9 patients with C-GBM tumors had PD-1^high^ CD4 T cell infiltration and 8/9 patients had PD-1^high^ CD8 infiltration whereas 2/10 patients with NC-GBM tumors had PD-1^high^ CD4 T cell infiltration and 3/10 NC-GBM patients had PD-1^high^ CD8 T cell infiltration. Further, mmRAPID identified a 2-fold increase in CD32 expression on CD8 T cells (mmRAPID Cluster 24) with a 2.6-fold decrease in patient survival (**Figure 5d**).

**Figure 5:**
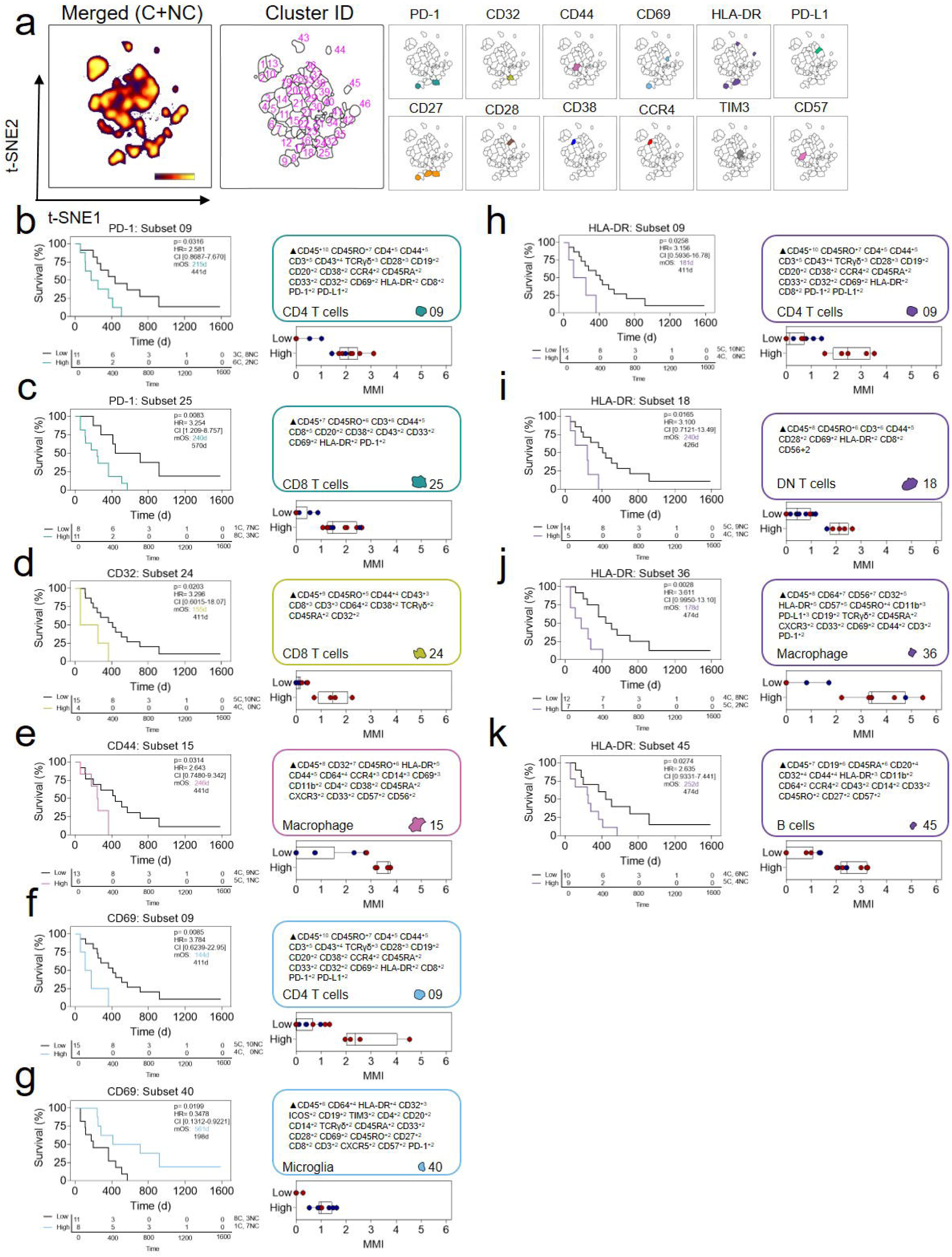
Immune receptor expression correlates with patient outcomes. **a**) T-SNE plots representing CD45+ cells pooled from n=19 GBM patients and enumeration of FlowSOM clusters on the t-SNE axes. Colored clusters indicate clusters identified by mmRAPID for which expression of the indicated markers stratify patient outcomes. **b-k)** Kaplan-Meier curves indicating overall survival in immune subsets stratifying patient outcome identified by mmRAPID analysis. Calculated MEM labels identified key features of stratifying immune subsets. Box and whisker diagrams plot the arcsinh transformed median mass intensity for PD-1 expression **(b, c)**, CD32 **(d)**, CD44 **(e)**, CD69 **(f, g)**, and HLA-DR **(h, i, j, k)** in stratified high and low groups.

Consistent with Citrus identification of elevated expression of CD44 on C-GBM infiltrative macrophages, mmRAPID found that 3.5-fold elevated expression of CD44 on macrophages (mmRAPID Cluster 15) correlated with 2-fold worse survival. The majority of patients (5/6) with CD44^high^ macrophages had C-GBM tumors, whereas 9/13 patients with CD44^low^ macrophages had NC-GBM tumors (**Figure 5e**). Elevated expression of CD69 on CD4 T cells (mmRAPID Cluster 9) correlated with poor outcome and lateral ventricle contact, but elevated CD69 expression on microglia (mmRAPID Cluster 40) was associated with more favorable outcomes and NC-GBM tumors (**Figure 5g**). Moreover, mmRAPID identified elevated HLA-DR expression on CD4 T cells, DNT cells, peripheral MDMs, and B cells as predicative of poor outcome and ventricle contact (**Figure 5h-k**). Surprisingly, PD-L1 expression, a feature of NC-GBM enriched microglia (**Figure 2**), correlated with an approximate 3-fold difference in survival outcomes and favored NC-GBM patients (**Supplementary Figure 8a**). Consistent with increased frequencies of CD27^+^ T cells in C-GBM tumors, we found that elevated CD27 expression on CD4, DNT, and CD8 T cells correlated with worse prognosis and favored patients with C-GBM tumors (**Supplementary Figure 8b-d**). While CD28 expression on microglia correlated with more favorable outcomes and NC-GBM tumors (**Supplementary Figure 8e**), CD38, CCR4, TIM3, and CD57 were associated with worse outcomes (**Supplementary Figure 8g-i**).

Taken together, these data identify a statistical association between immunoreceptor expression, lateral ventricle contact, and patient outcome as identified by two contrasting machine learning tools. Critically, elevated receptor expression tied to contact status and patient outcome identified by mmRAPID was consistent with Citrus identification of enriched immune populations in C-GBM and NC-GBM tumors. Co-expression of multiple predictive markers within the same immune subset suggests that key immune populations (e.g., PD-1^+^CD27^+^ T cells or HLA-DR^high^CD44^+^ MDMs) may be critical drivers of immunity within the ventricle tumor microenvironment. Paradoxical co-enrichment of activation markers (CD27, CD69, HLA-DR) as well as inhibitory receptors (PD-1) on the same subsets correlating with outcome suggested that these immune cells may be hyper-activated, may possess some functional signaling capacity.

### 6. Increased signaling capacity in the lateral ventricle tumor microenvironment

To address the signaling capacity of GBM infiltrating immune cells, we first used mass cytometry to assess the basal levels of 12 phosphoprotein readouts in immune infiltrates from 10 patients in our cohort (5 C-GBM, 5 NC-GBM) or healthy PBMC (**Figure 6a**, **Supplementary Table 1 and 4**). Basal levels of phosphorylated STAT1 (p-STAT1) were found only in myeloid cells from PBMC or GBM samples; however, basal levels of p-STAT3 were elevated in CD4 T cells, Tregs, B cells, and myeloid cells in GBM tumors compared to healthy donor PBMCs. Elevated basal STAT3 phosphorylation in myeloid cells in particular is consistent with infiltration of suppressive M2-like macrophages previously described (reviewed in (20)). Further, p-STAT5 levels were slightly increased in myeloid cells in the GBM tumor microenvironment. Ribosomal protein S6 (S6) and nuclear factor κB (NF-κB) were phosphorylated at baseline in T and B cells, and were highly S6 and NFκB phosphorylation, suggesting that while NK cells infiltrating GBM tumors may be functionally impaired, T, B, and myeloid populations appear to be functionally competent.

**Figure 6:**
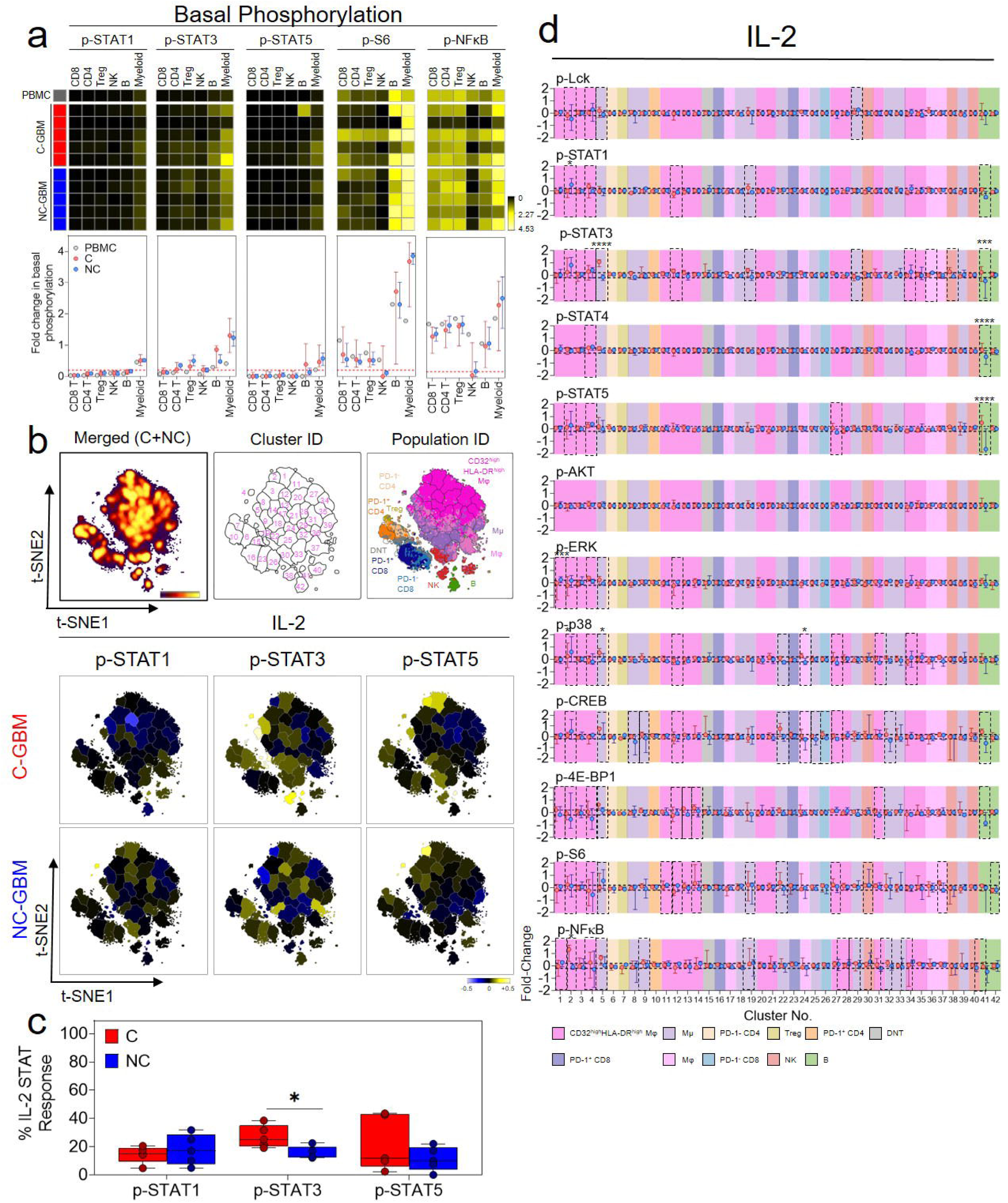
Immune cells infiltrating GBM tumors are functional and responsive to cytokine stimulation. **a)** Heatmaps indicating the arcsinh fold transformed median intensity values of each indicated phosphoprotein within each manually gated immune subset in healthy donor PBMC (grey), C-GBM tumors (red), or NC-GBM tumors (blue). **b)** Representative t-SNE plot indicating the density of CD45+ leukocytes (left), enumerated FlowSOM clusters (middle) and overlay of expert-gated immune populations onto the clustered t-SNE axes (right). Representative heatmaps on the t-SNE axes indicate the cluster-specific median arcsinh-fold change of the indicated phosphoprotein under IL-2 stimulation conditions compared to basal phosphorylation. **c)** Box and whisker plots indicating the proportion of clusters in C-GBM or NC-GBM immune infiltrates surpassing the phospho-signaling threshold (>0.2 arcsinh fold change) in response to IL-2 stimulation. **d)** Graphs indicating the arcsinh transformed phosphoprotein median intensities for each phosphoprotein in each of 42 IL-2 stimulated immune clusters. Graphs indicate the median ± IQR. Black horizontal lines in d indicate signaling threshold for each arcsinh transformed value. Black dotted boxes in d indicate clusters for which the arcsinh transformed median mass intensity surpassed the signaling threshold (±0.2-fold change) in either the C-GBM or NC-GBM cohort). * = p<0.05, ** = p<0.01, *** = p<0.001, **** = p <0.0001.

Given the co-enrichment of immune activation and inhibitory receptors and correlations with worse survival seen in C-GBM, we next assessed the ability of leukocytes in C-GBM and NC-GBM tumors to respond to cytokine stimulation. Bulk tumor samples were stimulated with cytokines with defined roles in regulating antitumor immunity including interleukin 2 (IL-2), IL-6, and interferon alpha (IFN α). FlowSOM clustering on t-SNE axes compared protein phosphorylation levels to basal signaling states in each of 42 defined immune subsets (**Figure 6b**). Quantification of the signaling responses for each of the 12 phosphoproteins within each cluster then identified which signaling networks were responsive to cytokine stimulation. Stimulation with IL-2 led to differential STAT phosphorylation within distinct immune subsets in C-GBM and NC-GBM tumors. For instance, STAT1 phosphorylation was highly refractory to IL-2 stimulation in C-GBM immune infiltrates, including Cluster 12 (a population of CD32^+^HLA-DR^high^ MDMs [see **Figure 4**]). Conversely, Cluster 2 (CD32^+^HLA-DR^+^ MDMs) exhibited phosphorylated STAT1 in response to IL-2 in NC-GBM tumors but not C-GBM tumors. Furthermore, IL-2 induced STAT3 phosphorylation in microglia (Clusters 5 and 19), and NK cells (Cluster 38) and STAT5 phosphorylation in CD32^+^HLA-DR^high^ MDMs (Clusters 2 and 4) and B cells (Cluster 41) in C-GBM tumors. Immune infiltrates in NC-GBM tumors, however, exhibited a different pattern of IL-2 mediated STAT3 phosphorylation, as Cluster 8 (microglia) and Cluster 38 (NK cells) failed to phosphorylate STAT3. Similar to C-GBM, Cluster 2 and 4 were responsive to IL-2 induced STAT5 phosphorylation, whereas Cluster 41 (B cells) was refractory to STAT5 phosphorylation. The proportion of clusters responding to IL-2 through STAT signaling revealed that immune infiltrates in C-GBM tumors were more significantly responsive to IL-2 through STAT3 (25% of clusters) than NC-GBM infiltrates (12.5% of clusters) (**Figure 6c**). Looking at the entire signaling profile, IL-2 induced several signaling cascades leading from membrane proximal signaling (LCK) to nuclear signaling (NF-κB) in several immune clusters (e.g. Cluster 2, 4, and 5) while other subsets (e.g. Cluster 41) appeared to be reciprocally impacted by IL-2 stimulation through multiple networks (**Figure 6d**).

Analogous to IL-2, IL-6 stimulation induced differential phospho-signaling networks in C-GBM and NC-GBM immune infiltrates (**Supplementary Figure 9a, b**). STAT3 phosphorylation was induced in CD32^+^HLADR^+^ MDMs (Clusters 2 and 21), microglia (Cluster 5), DNT cells (Cluster 15) and B cells (Cluster 41) infiltrating C-GBM, however these subsets failed to elicit a response to IL-6 in NC-GBM. (**Supplementary Figure 9a, b**). Interestingly, while IL-6 failed to induce STAT3 phosphorylation in Cluster 2 macrophages in NC-GBM patients, STAT5 appeared to be preferentially phosphorylated in this subset. In fact, IL-6 preferentially induced STAT3 phosphorylation across all C-GBM immune subsets, while STAT5 was favored in NC-GBM subsets (**Supplementary Figure 9c**), highlighting that inflammatory stimuli may have different immunomodulatory effects depending on tumor proximity to the lateral ventricle.

Stimulation with IFNα further distinguished immune responses in C-GBM and NC-GBM tumors (**Supplementary Figure 10a**). Interestingly, STAT1 and STAT3 appeared to be phosphorylated in microglia infiltrating C-GBM (Cluster 5) but not NC-GBM. In fact, a STAT1/STAT3-ERK-p38-CREB circuit appeared to be active in this population in C-GBM but not NC-GBM. Moreover, IFNα induced similar inflammatory circuits in CD32^+^HLA-DR^high^ macrophages in either C-GBM or NC-GBM. For example, STAT1, 3, and 4 were induced within Cluster 2 from both cohorts; however, STAT5 was preferentially induced in NC-GBM Clusters 2 and 5 as well as was an AKT-ERK-p38-CREB circuit specifically in Cluster 2 from NC-GBM patients (**Supplementary Figure 10b**), suggesting an active inflammatory response to IFNα in this subset. Similar to IL-2 and IL-6, IFNα stimulation appeared to favor STAT3 phosphorylation over STAT1 in C-GBM patients, as 19% of clusters demonstrated STAT3 responses in C-GBM, whereas only 10% of clusters demonstrated STAT1 phosphorylation, further supporting a role for STAT3 in immune regulation in C-GBM tumors.

Together, these data indicate that, not only do immune cells differentially infiltrate ventricle contacting and non-contacting tumors, but the inflammatory milieu within C-GBM may rewire immune signaling networks, drastically altering immune responsiveness to external stimuli. Strikingly, STAT3 phosphorylation appeared to drive much of the cytokine responsiveness in C-GBM tumors regardless of the cytokine stimulation or expected activation of canonical signaling pathways, consistent with our hypothesis that C-GBM tumors possess a distinct STAT3-driven immunosuppressive microenvironment.

## Discussion

Significant improvements in molecular and histologic characterization have increased our understanding of neurologic tumors, including glioblastoma. Unfortunately, improved tumor classification has not translated into clinical therapeutics that meaningfully impact patient outcomes. Work over the past decade has focused on tumor-specific characterization and targeting, and only recently has a large impetus been placed on understanding stromal factors that drive gliomagenesis and therapeutic resistance. The immune microenvironment, in particular, constitutes a critical part of tumor lesions and plays a crucial role in regulating tumorigenesis in peripheral and intracranial tumors (21, 22). While immune-targeted drugs have proven effective at generating antitumor immunity towards peripheral solid tumors and in some cases mediate complete tumor regression, the same efficacy has not been demonstrated in neurologic tumors (NCT02017717, NCT02667587) (23, 24). Early work using immunotherapeutic strategies has, in fact, demonstrated some capacity to elicit antitumor immunity, yet these approaches have not improved GBM outcomes (25–27), highlighting a need for novel, straightforward approaches, such as MRI-guided regional tumor position, to identify appropriate immune targets and patient cohorts to optimize therapeutic benefit.

Here we describe the immune microenvironment in glioblastomas based on regional tumor position, identifying immunomodulatory mechanisms in GBM tumors presenting with radiographic contact with the walls of the lateral ventricle. Complementary machine learning approaches provided supervised and unsupervised approaches that identified phenotypic and functional immune profiles associated with ventricle contact status and patient outcome. Namely, the abundance of lymphocytes and tissue resident microglia were increased in NC-GBM tumors, whereas C-GBM tumors were enriched in anti-inflammatory CD32^+^CD44^+^HLA-DR^high^ M2-like MDMs and exhausted PD-1^+^TIGIT^+^T cells. These observations suggest that C-GBM tumors, and potentially the periventricular space itself, are highly immunosuppressive environments.

Several lymphocyte subsets were enriched in NC-GBM tumors including γδ T cells and CD8^-^CD4^-^ DNT cells, and CD8 T cells, NK cells, and B cells which correlated with improved patient survival. Both γδ T cells and DNT cells can generate neuro-inflammatory responses during brain pathology, and are associated with antitumor cytotoxicity in tumors. Although T_EMRA_ cells are terminally differentiated, they possess increased cytotoxicity and tumor-killing capacity (18, 30), suggesting that T_EMRA_ cells may contribute to tumor control in NC-GBM tumors. Moreover, NK cells infiltrating NC-GBM tumors bore a CD56^low^ phenotype associated with potent toxicity, whereas CD56^high^ NK cells infiltrating C-GBM tumors suggest a regulatory capacity (reviewed in (31, 32)). Infiltration of multiple lymphocyte subsets with cytotoxic potential in NC-GBM suggests a more effective antitumor immune response in the microenvironment, and thus contribute to longer survival. Furthermore, B cells were enriched in NC-GBM. Recent reports have correlated development of B cell enriched tertiary lymphoid structures (TLSs) in tumors with improved patient survival and responsiveness to immunotherapy (33–35); however, the precise role of B cells in NC-GBM and their ability to support TLSs in NC-GBM tumors remains to be seen.

In contrast to NC-GBM tumors, lymphoid infiltration into C-GBM was reduced, and lymphocytes in C-GBM tumors bore a hyper-activated, exhausted phenotype characterized by increased expression of CD32, CD69, HLA-DR, CD27, and PD-1 on C-GBM T cells that correlated with worse prognosis. An increased frequency of PD-1^+^ T cells infiltrating C-GBM co-expressed a second checkpoint receptor, T cell immunoreceptor with Ig and ITIM domains (TIGIT). While the precise mechanism of TIGIT-mediated immune suppression is unclear, evidence suggests that TIGIT serves a compensatory role to PD-1-mediated inhibition, eliciting resistance to PD-1 targeted checkpoint blockade therapy (36). Moreover, TIGIT expression is associated with elevated T cell activation, immune exhaustion, and responsiveness to checkpoint blockade (37–39). Functionally, T cells in both C-GBM and NC-GBM expressed limited Granzyme B and Ki-67, suggesting a lack of effector function and proliferative capacity consistent with immune exhaustion. In contrast to T_EMRA_ infiltration in NC-GBM, CD45RO^+^CCR7^+^ central memory T cell phenotypes (T_CM_) predominated in C-GBM tumors. In fact, several T cell markers correlating with patient outcome (CD32, CD69, HLA-DR, CD27) are associated with T cell memory phenotypes. Whether T_EMRA_ and T_CM_ differentially infiltrate NC-GBM and C-GBM tumors, respectively, or are polarized by microenvironmental factors, and their functional consequences upon arrival remains to be determined. Although, increased frequencies of multiple CCR7^+^ lymphocyte populations in C-GBM tumors suggest that a CCR7-CCL19/CCL21 axis may recruit T_CM_ into the periventricular niche. Importantly, T cells in both C-GBM and NC-GBM tumors exhibited basal phosphorylation of both ribosomal protein S6 and NF-κB, indicating incomplete exhaustion and retention of some degree of functionality. While lymphocytes in both C-GBM and NC-GBM showed impaired responses to inflammatory cytokines (**Figure 7**), (arguing for exhaustion), lymphocytes in C-GBM tumors demonstrated moderate responses to inflammatory cytokines, particularly through CREB. It remains to be seen what effects other cytokines and soluble factors may have on T cell signaling, recruitment, and survival in C-GBM and NC-GBM.

**Figure 7:**
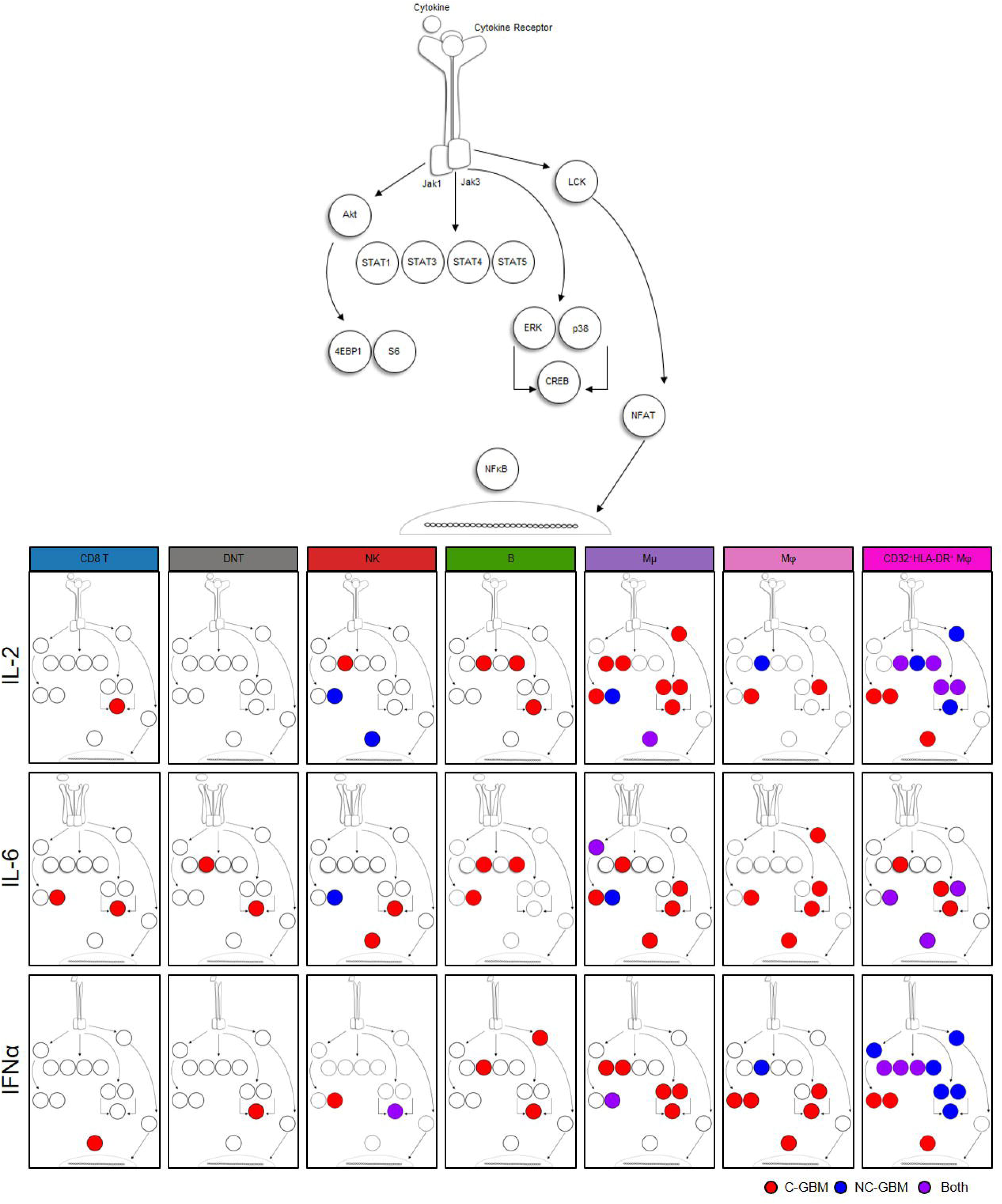
Model of cell signaling networks in GBM immune infiltrates. Graphical representation of immune cell signaling networks. For each cytokine stimulation condition implemented (rows) and each cell population of interest (columns), an aggregate signaling diagram was generated. Signaling nodes in red indicate active signaling responses to the indicated cytokine stimuli in C-GBM tumors and blue nodes indicate active signaling responses in NC-GBM. Purple nodes indicate protein phosphorylation in response to stimuli in both patient cohorts.

Tumor contact with the lateral ventricle also impacted phagocytic populations in the tumor microenvironment. A subset of CD45^low^CD11b^low/-^HLA-DR^low^CD14^-^ microglia was enriched in NC-GBM tumors and corresponded to more favorable outcomes, however, no distinct phenotype emerged to illuminate their functional capacity. Microglia in NC-GBM tumors demonstrated limited responsiveness to inflammatory cytokines with S6 phosphorylation remaining an active signaling component (**Figure 7**). A key role for microglia lies in maintenance of tissue homeostasis, in large part through phagocytosis of cellular debris and tissue pruning (reviewed in (40). The phagocytic capabilities of NC-GBM resident microglia and ability to present antigen is unknown, but may support infiltrating lymphocytes in mediating tumor control. In contrast to NC-GBM tumors, two populations of peripheral CD45^high^CD11b^+^CD14^+^ MDMs were enriched in C-GBM tumors. These populations were distinguished by expression of CD32, CD44, HLA-DR, CD69, and chemokine receptors CXCR3 and CCR4. Along with elevated frequencies of CCR7+ lymphocytes, these findings further support a role for chemotactic factors in recruiting leukocyte populations into the niche.

Functionally, C-GBM infiltrating MDMs expressed CD163 and CD209 (DC-SIGN), indicative of M2-like anti-inflammatory macrophages in the tumor microenvironment (41). MDMs also exhibited high basal expression of phosphorylated STAT3, consistent with M2-like polarization. Interestingly, CD32^+^HLA-DR^+^ macrophages showed greater responsiveness to cytokine stimulation than their CD32^-^HLA-DR^-^ counterparts (**Figure 7**). Upon stimulation, these MDMs utilized different signaling networks in C-GBM and NC-GBM tumors. For example, CD32^+^HLA-DR^+^ macrophages signaled through a STAT 3/4/5-ERK- p38-CREB axis in NC-GBM tumors in response to IL-2 stimulation. While this network was active in C-GBM MDMs, 4-EBP1, S6, and NF-κB were also phosphorylated, suggesting some degree of differential macrophage signaling in these regional tumor classes. Similarly, C-GBM MDMs were more responsive to IL-6 stimulation, particularly through STAT3 in C-GBM compared to NC-GBM. Perhaps most striking, CD32^+^HLA-DR^+^ MDMs were highly responsive to IFNα stimulation. While all STAT proteins assessed were phosphorylated following IFNα stimulation in C-GBM and NC-GBM MDMs, NC-GBM MDMs favored an inflammatory ERK-p38-CREB signaling axis, while C-GBM MDMs favored 4-EBP1, S6, and NF-κB signaling, further supporting a role for the tumor environment in mediating differential myeloid signaling responses. Recent reports have demonstrated conflicting roles for IFNα signaling in mediating pro-or antitumor responses depending on the chronicity of IFN exposure (see (42)). Our results support a hypothesis that long-term IFN exposure in the ventricular space may negatively impact antitumor immunity, whereas acute inflammatory IFN signaling along with increased lymphocyte infiltrate in NC-GBM supports antitumor immunity and prolonged overall survival in patients with NC-GBM tumors. Importantly, STAT3 phosphorylation played a role in C-GBM immune infiltrates regardless of stimulation condition. This points to STAT3 as a critical, targetable, driver of antitumor immunity in the ventricular space, and suggests the stoichiometry of STAT3 may enforce a regulatory immune signaling axis; however, the ability of STATs to form heterodimeric complexes in response to inflammatory cues in brain tumors, and the resulting functional consequences, is poorly understood.

We assessed signaling responses to three inflammatory cytokines that impact antitumor immunity to peripheral solid tumors; however, a complex milieu of cytokines, soluble mediators, and unique parenchymal factors within the brain may further influence antitumor immunity in the brain. Identifying the cellular source--be it tumor-, stem cell-, ependyma-, or immune-derived—and the dynamic interplay between these cell subsets within the microenvironment in mediating antitumor immunity in C- and NC-GBM tumors will be critical to advance our understanding of neuro-oncology and develop novel therapeutics.

This work highlights potential immunotherapeutic targeting strategies for GBM patients based on MRI-guided tumor proximity to the lateral ventricle. Future studies will be needed to determine whether patients with NC-GBM or C-GBM tumors may be more amenable to drug combinations targeting lymphocyte of MDM populations respectively. Immuno-oncology agents currently approved or in clinical trials may afford the most immediate benefit, particularly for patients with C-GBM tumors; including agents targeting PD-1 (nivolumab, pembrolizumab), TIGIT (tiragolumab), CD27 (varlilumab), or STAT3 (WP-1066). It remains to be seen, however, which immunotherapeutic combinations will lead to improved outcomes in GBM patients.

The tumor immune microenvironment is heavily influenced by tumor tissue-of-origin, particularly in the brain. For instance, brain metastases possess distinct immune microenvironments dependent on tissue of origin (43). Here, we demonstrate that regional position of primary brain lesions, visualized as MRI-guided contact with the lateral ventricle, influenced antitumor immunity in the brain and will be germane to clinical decision making, particularly in patient selection and therapeutic options available to patients with C-GBM vs NC-GBM.

## Methods

### Human Specimens

Freshly resected glioblastoma tumor tissues were collected from the Department of Neurosurgery at Vanderbilt University Medical Center between 2014 and 2018 as previously described (13,44,45). Non-tumor bearing brain tissue was collected from temporal lobectomy as standard-of-care treatment for epilepsy from the Veterans’ Affairs Medical Center affiliated with Vanderbilt University. Glioblastomas with isocitrate dehydrogenase (IDH) mutations confirmed by standard pathological diagnosis were excluded from this study. All samples were collected with patient informed consent in compliance with the Vanderbilt Institutional Review Board (IRBs #030372, #131870, #181970), and in accordance with the Declaration of Helsinki. Samples were de-identified prior to tissue processing. All patients were adults (age range 40-80 years old) at the time of tumor resection. Extent of resection was classified as gross total resection (GTR) or subtotal resection (STR) independently by a neurosurgeon and a neuro-radiologist. Gross total resection was defined as no significant residual tumor enhancement upon gadolinium enhanced magnetic resonance imaging (MRI) of the brain 24 hours’ post-surgery. Tumor contact with the lateral ventricle was confirmed upon inspection of MRI and verified by a neurosurgeon and neuro-radiologist. All patients were considered for postoperative chemotherapy (temozolomide) and radiation according to standard-of-care. Methylation of the O^6^-methylguanine-DNA methyltransferase (MGMT) promoter was determined by pyrosequencing (Cancer Genetics Inc, Los Angeles CA, USA). *IDH* mutation status was confirmed using polymerase chain reaction (PCR). Patient follow-up extended to October 2019, noting time to first radiographic progression or tumor recurrence as assessed by a neuro-oncologist and neuro-radiologist or time to patient’s death. All deaths were deemed related to tumor progression. Median overall survival (mOS) of the patient cohort presented herein was 366 days. A total of 25 glioblastoma patients from both male and female subjects were included in the present study. A complete list of clinical characteristics can be found in (**Supplementary Table 1**).

Peripheral blood mononuclear cells (PBMC) were collected from healthy volunteers with written informed consent under IRB protocol #131311 and in accordance with the Declaration of Helsinki. Samples were de-identified prior to processing, and no other information was obtained from healthy individuals.

### Tissue Collection and Processing

Fresh tumor tissue was obtained directly from the operating room at Vanderbilt University Medical Center within one hour of resection. Tissues were processed and dissected into single-cell suspensions as previously reported (44, 45). Briefly, samples were resuspended in neural stem cell media: DMEM-F12 supplemented with glutamine (Glutamax, Thermo Fisher), 1M HEPES (Gibco), a hormone cocktail consisting of 30% glucose (Fisher), 7.5% sodium bicarbonate (Sigma Aldrich), apotransferrin, insulin, Putrescine Solution, 200 μM Progesterone, 3 mM Sodium Selenite (Sigma Aldrich), and gentamycin (Fisher). Tissues were minced with razor blades to an approximate diameter of 1 mm before enzymatic digestion for 1 hour with collagenase IV (1 ug/ul) and DNAse I (0.25 ug/ul) at 37°C and 5% CO_2_ with steady shaking. Suspensions were then washed and triturated before filtration twice through 70 um and 40 um filters. Cell pellets were resuspended in ACK lysis buffer to remove red blood cells (Invitrogen). Cells were washed and resuspended in neural stem cell media supplemented with bovine serum albumin (Sigma Aldrich), heparin, recombinant human FGF (25ug/mL; Stem Cell Technologies), and recombinant human EGF (10 ug/mL; Stem Cell Technologies) in 10% DMSO before long-term cryopreservation (1X10^7^ cells/mL) in liquid nitrogen.

Healthy donor blood was collected by venipuncture into heparinized collection tubes (Becton Dickinson; 100 mL/donor). Whole blood was diluted 1:4 with PBS before being overlaid onto a Ficoll-Paque Plus density gradient (GE Lifesciences) Blood was then centrifuged at 400g for 30 minutes without a brake. Buffy coats were isolated, washed with PBS, and centrifuged at 500g for 10 minutes. Cell pellets were then resuspended in ACK lysis buffer for 5 minutes, washed, and cryopreserved at 1X10^7^ cells/mL in liquid nitrogen in 10% DMSO in FBS.

### Metal-isotope Tagged Antibodies

All antibodies used for mass cytometry analysis are listed in **Supplementary Table S2, Supplementary Table S3, and Supplementary Table S4**. Pre-conjugated antibodies to metal isotopes were purchased from Fluidigm or from commercial suppliers in purified form and conjugated in house using the Maxpar X8 chelating polymer kit (Fluidigm) according to the manufacturer’s instructions.

### Cell Preparation and Mass Cytometry Acquisition

Cryopreserved samples were rapidly thawed in a 37°C water bath and resuspended in complete RPMI supplemented with 10% FBS and 50 units/mL of penicillin–streptomycin (Thermo Scientific HyClone). Cell suspensions were processed and stained as previously described (44, 46). Briefly, cells were washed once with serum free RPMI and subsequently stained with ^103^Rh Cell-ID Intercalator (Fluidigm) at a final concentration of 1 uM for 5 minutes at room temperature. Staining was quenched with complete RPMI before washing with PBS 1% BSA. Cells were resuspended in PBS/BSA and added to the appropriate antibody cocktail of cell-surface staining antibodies (**Supplementary Table S2, Supplementary Table S3, Supplementary Table S4**) and incubated at room temperature for 30 minutes. Samples were washed in 1% PBS/BSA before fixation in 1.6% paraformaldehyde for 10 minutes at room temperature. Cells were again washed in PBS and fixed in ice cold methanol with gentle vortexing before storage at -20°C. On the day of data collection, samples stored at -20°C were washed in PBS/BSA and resuspended in an antibody cocktail of intracellular stains (e.g., granzyme B, VISTA) for 30 minutes. Iridium Cell-ID Intercalator was added at a final concentration of 125 nM and incubated at room temperature for at least 30 minutes. Cells were then washed and resuspended in ultrapure deionized water, mixed with 10% EQ Four Element Calibration Beads (Fluidigm) and filtered through a 40 uM FACS filter tube before data collection. Data were collected on a Helios CyTOF 3.0 (Fluidigm). Quality control and tuning processes were performed following the guidelines for the daily instrument operation. Data were collected as FCS files.

### Cytokine Stimulation and Phospho-specific Cytometry

Phospho-specific mass cytometry was performed as previously described (46). Cryopreserved samples were thawed in a water bath as above. Cell pellets were resuspended in complete RPMI (10% FBS, penicillin-streptomycin) and rested at 37°C for 15 minutes. Cell suspensions were then washed in PBS and stained in 1 µM rhodium Cell-ID intercalator in PBS for 5 minutes. Cells were again washed in PBS/BSA and aliquoted equally into cytokine stimulation solutions. Briefly, these conditions included PBS (unstimulated), recombinant human interleukin 2 (20 ng/mL), recombinant human interleukin 6 (20 ng/mL), recombinant human interferon alpha (20 ng/mL) or hydrogen peroxide (10uM). Stimulation conditions were allowed to proceed for 15 minutes before immediate fixation in 1.6% PFA in order to halt phospho-protein dissociation. Samples were then washed in PBS/BSA, stained with a cocktail of cell-surface antibodies and fixed in ice cold methanol as above. On the day of collection, samples were stained with a cocktail of intracellular phospho-specific antibodies. Iridium intercalation, resuspension in EQ calibration beads, and sample collection proceeded as above.

### Data Preprocessing

Raw mass cytometry files were normalized using the MATLAB bead normalization tool. Files were then uploaded to the cloud-based analysis platform Cytobank. Before automated high-dimensional data analysis, the mass cytometry data were transformed with a cofactor of 5 using an inverse hyperbolic sine (arcsinh) function. Cell doublets were first excluded using Gaussian parameters (Center, Offset, Width, Residual) as reported (46). Intact cells were gated based on DNA content (^191^Ir and ^193^Ir). Dead cells were excluded based on rhodium intercalation. Immune subsets were then manually gated using biaxial gating strategies.

### Dimensionality Reduction and Automated Clustering

T-distributed stochastic neighbor embedding (t-SNE) analysis was performed on each individual patient sample using the Cytobank platform. To avoid down-sampling, all live CD45+ cells were included for each individual patient’s t-SNE analysis (range: 1,322-335,303 events) using all immune markers in the antibody panel to generate t-SNE maps, a perplexity of 30, theta of 0.5, and 10,000 iterations. Automated clustering of phenotypically distinct immune cell subsets was then performed for each individual patient using the Cytobank implementation of the FlowSOM clustering tool(9). Briefly, clustering analysis was performed using only the t-SNE1 and t-SNE2 channels, hierarchical consensus clustering, a cluster number of 196, and 10 iterations. Multiple FlowSOM analyses using different iterations of metaclusters (5-50) identified the optimal number of total metaclusters that minimized the variance of each immune marker in each metacluster as previously described (13). For patient-to-patient comparisons where indicated, an equal number of live CD45+ events were down-sampled from each patient file, concatenated together, and the number of FlowSOM clusters was optimized across all patients.

### Citrus Clustering

To identify immune populations that may be enriched in abundance in either lateral ventricle contacting or non-contacting glioblastomas, we employed the Cytobank implementation of the Citrus clustering algorithm (12, 47). Live CD45+ cells were equally downsampled from 19/25 patient files (478 events/patient) (**Supplementary Table S1**). Patient files included in the analysis were grouped by tumor contact with the lateral ventricle (9 contacting, 10 non-contacting). All immune markers in the Immune Phenotyping Panel (**Supplementary Table S2**) were used to cluster. A nearest shrunken centroid predictive analysis of microarrays (PAMR) model was used to predict enriched cell abundance within each group. The minimum cluster size was set to 5%, with 5 cross validation folds, and a false discovery rate (FDR) of 1%. To determine enriched immune marker expression, the Citrus algorithm was rerun investigating median marker expression. Sixteen immune markers delineating common lymphocyte and myeloid cell subsets were used to cluster events, while the arcsinh expression levels of 17 markers was explored (**Supplementary Table S2**). The maximum number of events per file was sampled. The same analysis settings used for abundance analysis was similarly used for median marker expression (PAMR analysis, 5% cluster size, 5 cross validation folds, FDR=1%). The most terminal clusters of differential abundance between the two cohorts of glioblastoma patients were exported into new experiments for further analysis. Biaxial gating was used to determine each cluster phenotype. Due to down-sampling in the Citrus analysis, and in order to determine from which immune populations each Citrus cluster was sampled, t-SNE analysis was performed on the total number of live CD45+ cells from each patient as well as each patient’s individual Citrus clusters. Cells from Citrus clusters were assigned to FlowSOM metaclusters based on 1) position in the overlaid viSNE map, and 2) phenotypic similarity 3) proportion of FlowSOM metaclusters. Further, both abundance of cells with an identified Citrus phenotype and expression of identified markers were validated independently in the entire 25-patient cohort.

### Marker Enrichment Modeling

The phenotypes of automatically clustered immune populations generated in Citrus and FlowSOM were identified using Marker Enrichment Modeling (MEM) (11, 48). Briefly, clusters generated from FlowSOM or Citrus were exported from Cytobank into R. Within R, MEM labels were generated using all immune markers included in either the immune phenotyping cytometry panel (**Supplementary Table S2**) or the immune checkpoint cytometry panel (**Supplementary Table S3**). As a modification of the original MEM script, herein we compared marker expression within each cluster to a statistical null reference point wherein the magnitude of the median expression value of the null set was defined as 0, and the interquartile range (IQR) was defined as the median IQR for all features in the MEM analysis (49). MEM values were then scaled from 0 (no expression) to 10 (high expression) relative to the reference point. Populations were then hierarchically clustered using the hclust package in R based on median marker expression, MEM value, or IQR. MEM labels for each population were confirmed by biaxial gating and used to derive an expert-guided population identity.

### Root Mean Square Deviation

To compare intra- and inter-patient phenotypic similarities between automatically clustered immune populations, MEM values were generated for each cluster as indicated. MEM labels were then used to compare each clustered population using the Root-mean-square-deviation (RMSD) calculation in the “MEM_RMSD” function included in the MEM package in R (https://github.com/cytolab/mem). Briefly, the “MEM_RMSD” function calculates the square root of the mean squared distance between every MEM value in common for a given pair of cell subsets. These values are then transformed and expressed as a percentage of the maximum RMSD in the analysis. Heatmaps of each hierarchically clustered population based on RMSD score and a matrix of RMSD values were exported from R. MEM labels were generated by concatenating all clusters within a branch of the RMSD hierarchal clustering tree, and an average MEM value for each marker was generated.

### Analysis of phospho-signaling

Analysis of immune cell phospho-signaling under different cytokine stimulation conditions was performed in Cytobank. Baseline phospho-signaling was compared in biaxially-gated populations. Changes in protein phosphorylation under stimulation conditions were normalized to basal signaling in unstimulated automatically clustered FlowSOM populations. Briefly, an equal number of CD45+ events (3523/patient) was first down-sampled from each patient’s stimulation condition file before t-SNE analysis as above. All immune phenotypic markers were used to generate the t-SNE plot, while phospho-protein readouts were excluded. FlowSOM clustering analysis was performed on the t-SNE axes and the number of FlowSOM clusters was optimized to minimize variance in marker expression. The median expression value for each phospho-protein readout from each stimulation condition was then compared to the median expression value from unstimulated cells using the median arcsinh transformation. A signaling threshold ≥ 0.2 or ≤ -0.2 in arcsinh transformed median phospho-protein expression over baseline was considered a cellular response to each stimulation condition.

### Survival Analysis

Correlations of immune subset abundance with overall and progression-free survival were performed using the Risk Assessment Population IDentification algorithm (RAPID) in R (13). Briefly, t-SNE analyses were performed by down-sampling live CD45+ events from each patient file and optimizing the number of FlowSOM clusters as described above. The frequency of cells from each patient within each cluster was then used to stratify patients into high and low cluster abundance based on the interquartile distribution of the subset across the entire cohort. A univariate Cox regression model was then used to estimate the hazard ratio of death and determine statistical significance using in the “survival” package in R. Overall survival (OS) was defined as time from surgical resection to death. Survival time was censored if, at last follow-up, the patient was known to be alive and had not had radiographic tumor progression. Differences in the survival curves were compared using the Cox univariate regression model, reporting a hazard ratio (HR) between the survival curves. Statistical significance α was set at 0.05 for all statistical analyses. RAPID analysis was performed on 10 different t-SNE analyses resampled from each patient. Clusters from each independent analysis that met an HR threshold >1 or <-1 and p-value <0.1 were isolated and RMSD was performed to determine the stability of clusters with similar phenotypes identified by RAPID.

A modified version of the RAPID algorithm (mmRAPID) was used to correlate immune receptor expression with patient outcome. Dimensionality reduction was performed using t-SNE on 16 of the clustering markers used in the Citrus median marker analysis (**Supplementary Table S2**). FlowSOM clustering was then performed on the t-SNE axes to minimize the variance in marker expression across all clusters as previously described. The arcsinh transformed median marker expression of 17 markers of interest was used to stratify patients into high and low expression based on the interquartile distribution of the arcsinh transformed values across the entire cohort for each cluster. A univariate Cox regression model was used to estimate the hazard ratio of death as was performed for immune abundance. A similar analysis correlating phospho-signaling to survival outcome was performed using the arcsinh transformed phospho-protein expression values normalized to the unstimulated baseline levels for each patient. Marker expression was validated by biaxial gating and generation of MEM labels for high and low groups.

### Statistical Analysis

Statistical analysis was performed in Cytobank as a part of advanced analyses, in R version 3.6.1, or Graphpad Prism version 8.4.3 where indicated. All analyses were graphed in the Graphpad Suite. Outlier analysis was performed before all statistical analyses using the ROUT method with Q=1%. Statistical analysis of 2 groups was performed using a two-sided student’s T test with Welch’s Correction. Analysis of 3 or more groups was performed where indicated using a one-way or two-way ANOVA with a Tukey or Sidak correction for multiple hypothesis testing, respectively. Immune subset enrichment was statistically determined using a Chi Squared or Fishers Exact Test where indicated. Statistical correlations of immune subset abundance were performed using a two-tailed Pearson Correlation. For all statistical tests unless otherwise indicated, *p* values of <0.05 were considered significant. Graphs show median ± IQR unless otherwise indicated.

### Data Availability

Datasets analyzed in this manuscript will be made available online, including at FlowRepository (50), for reviewers and at the time of publication. Transparent analysis scripts for datasets in this manuscript (first shown in **Figure 2** and **Figure 5**) will be publicly available on the CytoLab Github page (https://github.com/cytolab/GBM-IMM01) with open-source code and commented Rmarkdown analysis walkthroughs.

## Acknowledgements

We thank Vanderbilt’s Cancer and Immunology Core and Flow Cytometry Shared Resource facilities as well as all the surgeons, patients, and families that supported this work. Research was supported by the following funding resources: R01 NS096238 (R.A.I., J.M.I.), R01 CA226833 (J.M.I., S.M.L., T.B.), U01 AI125056 (J.M.I., S.M.B.), U54 CA217450 (J.M.I.), K00 CA212447 (T.B.), Burroughs Wellcome Fund 1018894 (A.M.M.), the Michael David Greene Brain Cancer Fund (R.A.I., J.M.I.), the Southeastern Brain Tumor Foundation (R.A.I., J.M.I.), and the Vanderbilt-Ingram Cancer Center (VICC, P30 CA68485).

## Author Contributions

T.B., R.A.I., and J.M.I. designed the study. T.B., M.J.H., J.S., N.L., and C.E.R. collected data. T.B. and S.M.L. developed data analysis scripts. T.B., S.M.L., R.A.I., and J.M.I. performed mass cytometry data analysis and interpretation. A.M.M. scored MRI images for tumor contact with the lateral ventricle and provided patients’ clinical characteristics. B.C.M. confirmed tissue pathology. L.B.C., R.C.T., and K.D.W. provided freshly resected tissue specimens. T.B., R.A.I., and J.M.I. wrote the manuscript. R.A.I. and J.M.I. provided financial support. All authors contributed to reviewing and editing the manuscript.

