## Supplementary Figures and Figure Legends for "Immune cell regulation in stem cell niche contacting glioblastomas"

### Supplementary Figure 1

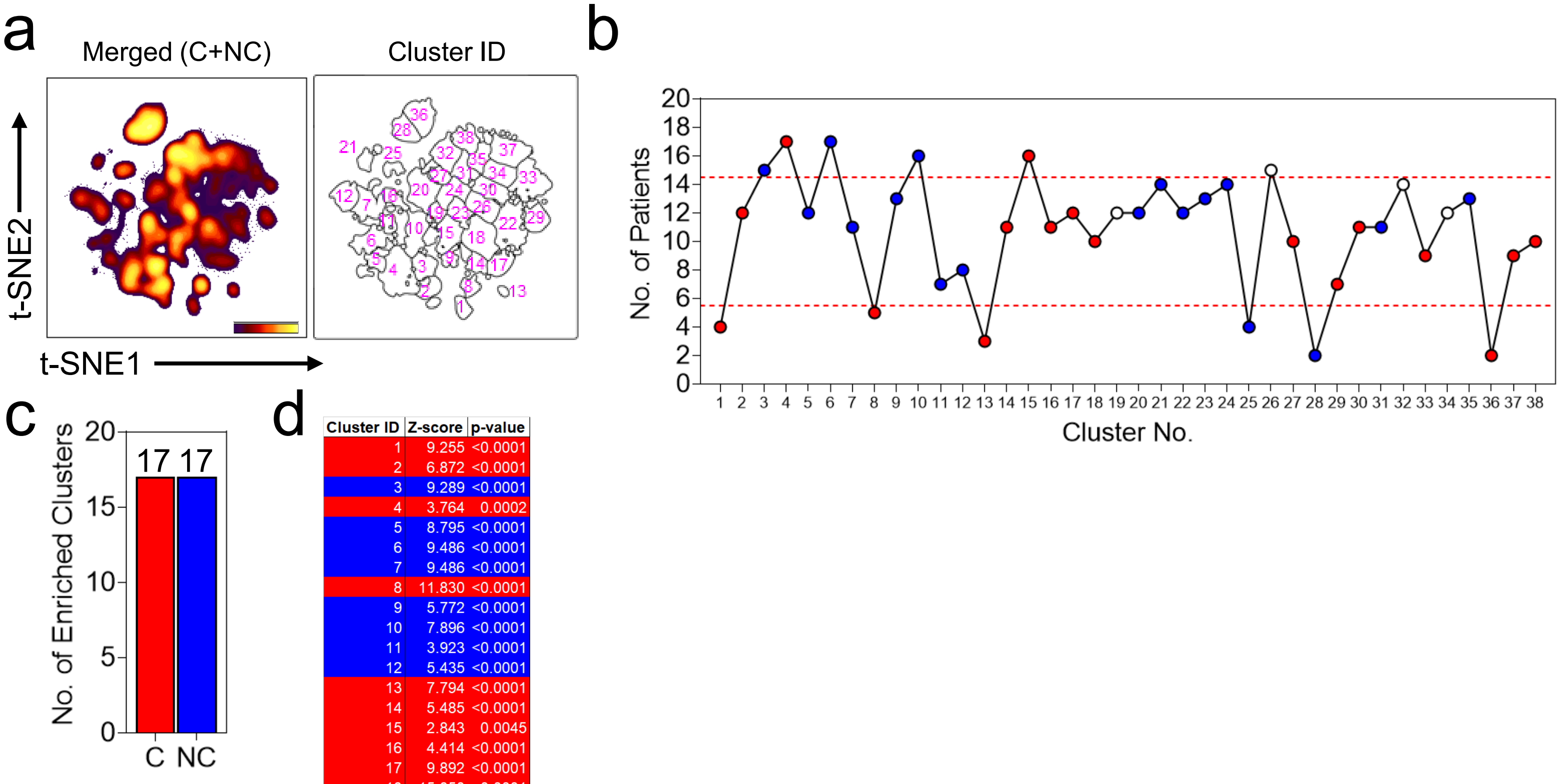


| Cluster ID | Z-score | p-value |
|---|---|---|
| 1 | 9.255 | <0.0001 |
| 2 | 6.872 | <0.0001 |
| 3 | 9.289 | <0.0001 |
| 4 | 3.764 | 0.0002 |
| 5 | 8.795 | <0.0001 |
| 6 | 9.486 | <0.0001 |
| 7 | 9.486 | <0.0001 |
| 8 | 11.830 | <0.0001 |
| 9 | 5.772 | <0.0001 |
| 10 | 7.896 | <0.0001 |
| 11 | 3.923 | <0.0001 |
| 12 | 5.435 | <0.0001 |
| 13 | 7.794 | <0.0001 |
| 14 | 5.485 | <0.0001 |
| 15 | 2.843 | 0.0045 |
| 16 | 4.414 | <0.0001 |
| 17 | 9.892 | <0.0001 |
| 18 | 15.850 | <0.0001 |
| 19 | 0.808 | 0.4192 |
| 20 | 13.110 | <0.0001 |
| 21 | 13.110 | <0.0001 |
| 22 | 10.740 | <0.0001 |
| 23 | 12.200 | <0.0001 |
| 24 | 5.451 | <0.0001 |
| 25 | 8.335 | <0.0001 |
| 26 | 1.895 | 0.0581 |
| 27 | 4.228 | <0.0001 |
| 28 | 8.386 | <0.0001 |
| 29 | 10.410 | <0.0001 |
| 30 | 2.535 | 0.0186 |
| 31 | 2.819 | 0.0048 |
| 32 | 0.810 | 0.4182 |
| 33 | 13.020 | <0.0001 |
| 34 | 1.081 | 0.2798 |
| 35 | 6.237 | <0.0001 |
| 36 | 7.917 | <0.0001 |
| 37 | 11.740 | <0.0001 |
| 38 | 8.058 | <0.0001 |

**Supplementary Figure 1: Computational determination of enriched immune populations in C-GBM and NC-GBM tumors.**

**a)** t-SNE plots indicating the density (left) and FlowSOM cluster enumeration of CD45+ cells pooled from 19/25 GBM patients. **b**) Graphical representation of the number of patients represented within the FlowSOM cluster. Dotted red lines indicate the 25th and 75th percentile respectively. Red dots indicate a statistical enrichment of C-GBM patients, blue dots indicate enrichment of NC-GBM patients in each cluster, and white dots indicate non-enriched clusters. **c)** Bar graph representing the number of total clusters statistically enriched in patients with C-GBM or NC-GBM tumors. **d)** Chart indicating the Z-scores and p-values from Chi-squared analysis to determine statistical enrichment of cells from C-GBM (red) or NC-GBM patients for each cluster.

Supplementary Figure 2

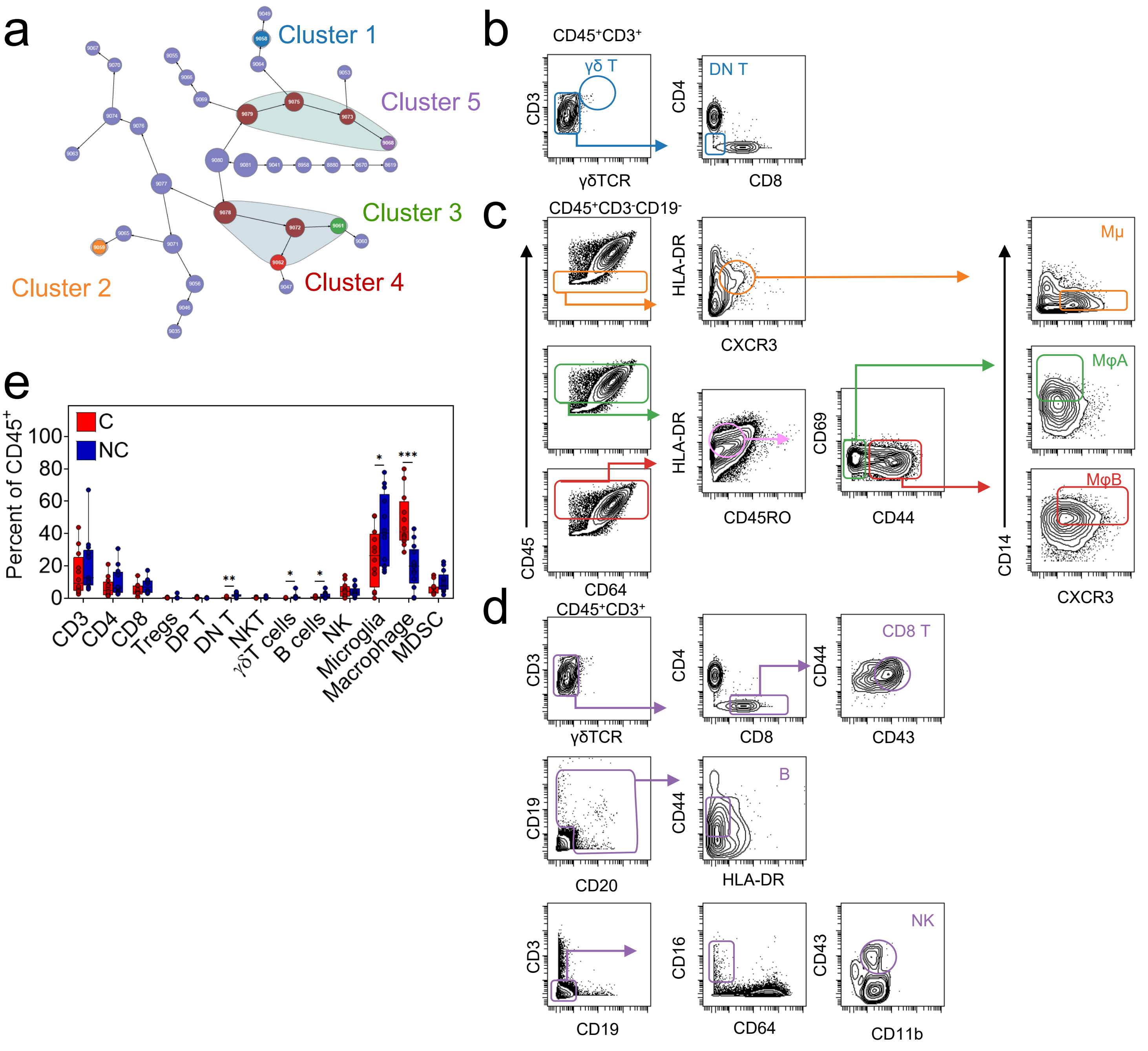


**Supplementary Figure 2: Identified Citrus clusters correspond to lymphoid and phagocytic immune subsets.**
**a)** Citrus clustergram of statistically enriched immune cell populations. Circled maroon nodes indicate clusters of differential abundance. Color coded nodes indicate terminal nodes used for further analysis. **b-d)** Representative expert-guided biaxial gates identifying immune cells subsets identified by Citrus in 6/25 patients not included in Citrus analysis. **e)** Box and whisker plots showing the frequency of each indicated expert-gated immune population as a percent of total CD45+ tumor infiltrating leukocytes. Gating strategies in b-d were defined by MEM labels and the most distinguishing features for each Citrus subset. Quantification in **e** represents manually gated populations from 25/25 patients in the cohort. * = p<0.05, ** = p<0.01, *** = p<0.001, **** = p <0.0001.

### Supplementary Figure 3

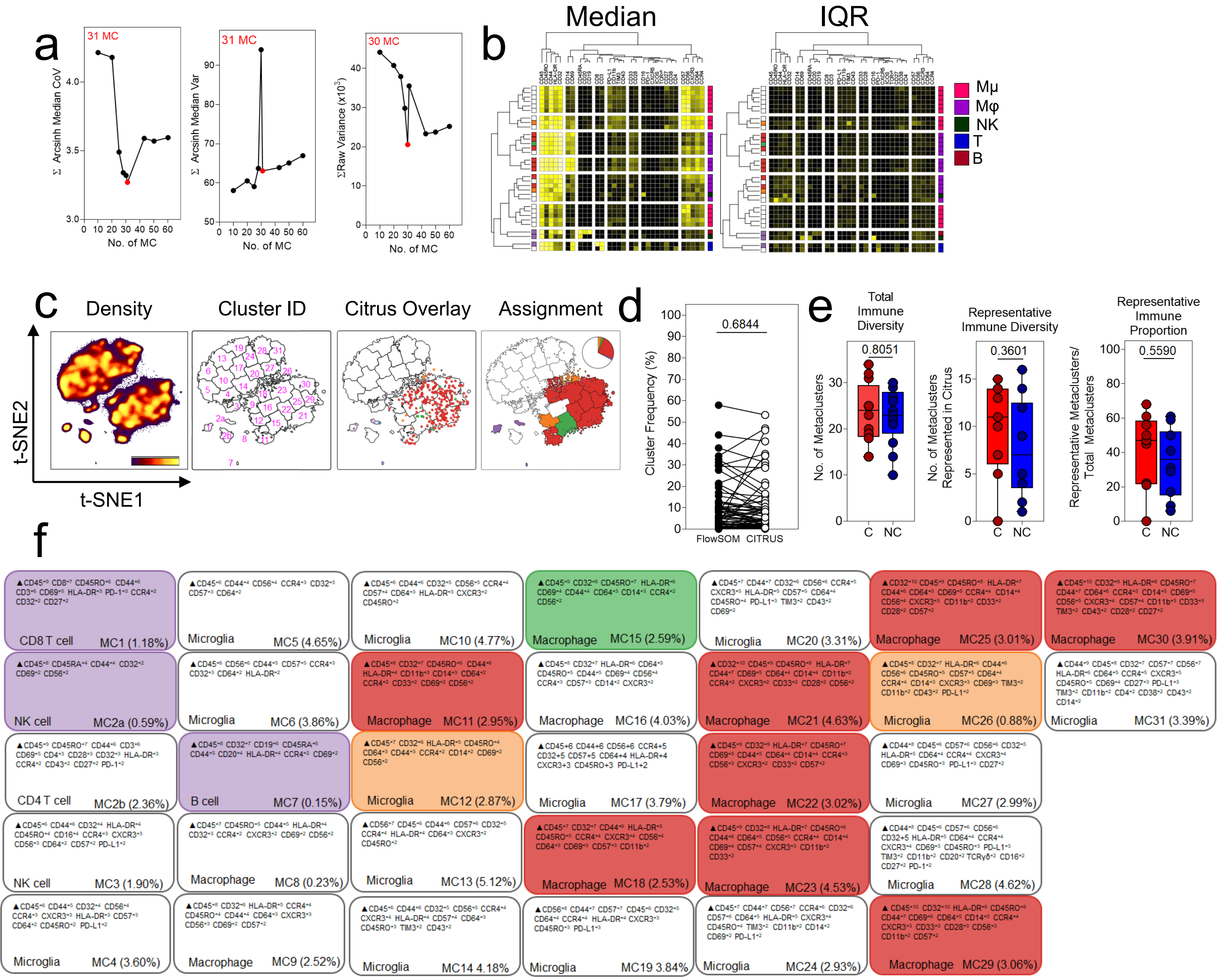


**Supplementary Figure 3: Optimization of FlowSOM clustering on t-SNE axes.**
**a)** Representative plots demonstrating minimization of the aggregate arcsinh transformed coefficients of variation (left), arcsinh transformed variances (middle), or raw variance (right) for each marker across multiple iterations of FlowSOM clustering. **b)** Representative heatmap of the arcsinh transformed median mass intensity (left) or interquartile range (IQR) (right) of each indicated marker across each FlowSOM cluster. Colored boxes in the left indicate Citrus identity. Colored boxes on the right indicate an expert-defined immune identify for each cluster. **c)** Representative assignment of patient-specific cells from Citrus clusters onto t-SNE axes including cell density (far left), enumeration of FlowSOM clusters on the t-SNE axes (mid left), overlay of Citrus populations on to the t-SNE plot (mid right) and assignment of Citrus populations for FlowSOM clusters from which they putatively were down sampled **d)** Comparison of the frequency of each Citrus cluster identified in each patient to the frequency of the FlowSOM cluster when accurately assigned on the t-SNE space. **e)** The number of optimized FlowSOM clusters per patient (left), total number of FlowSOM clusters containing cells from Citrus analysis (mid), and proportion of FlowSOM metaclusters containing Citrus cells was calculated. **f)** Representative MEM labels generated for each patient-specific FlowSOM cluster. Clusters were assigned labels based on marker expression and were color-coded to indicate clusters derived from Citrus. The frequency of each cluster is also indicated.

Supplementary Figure 4

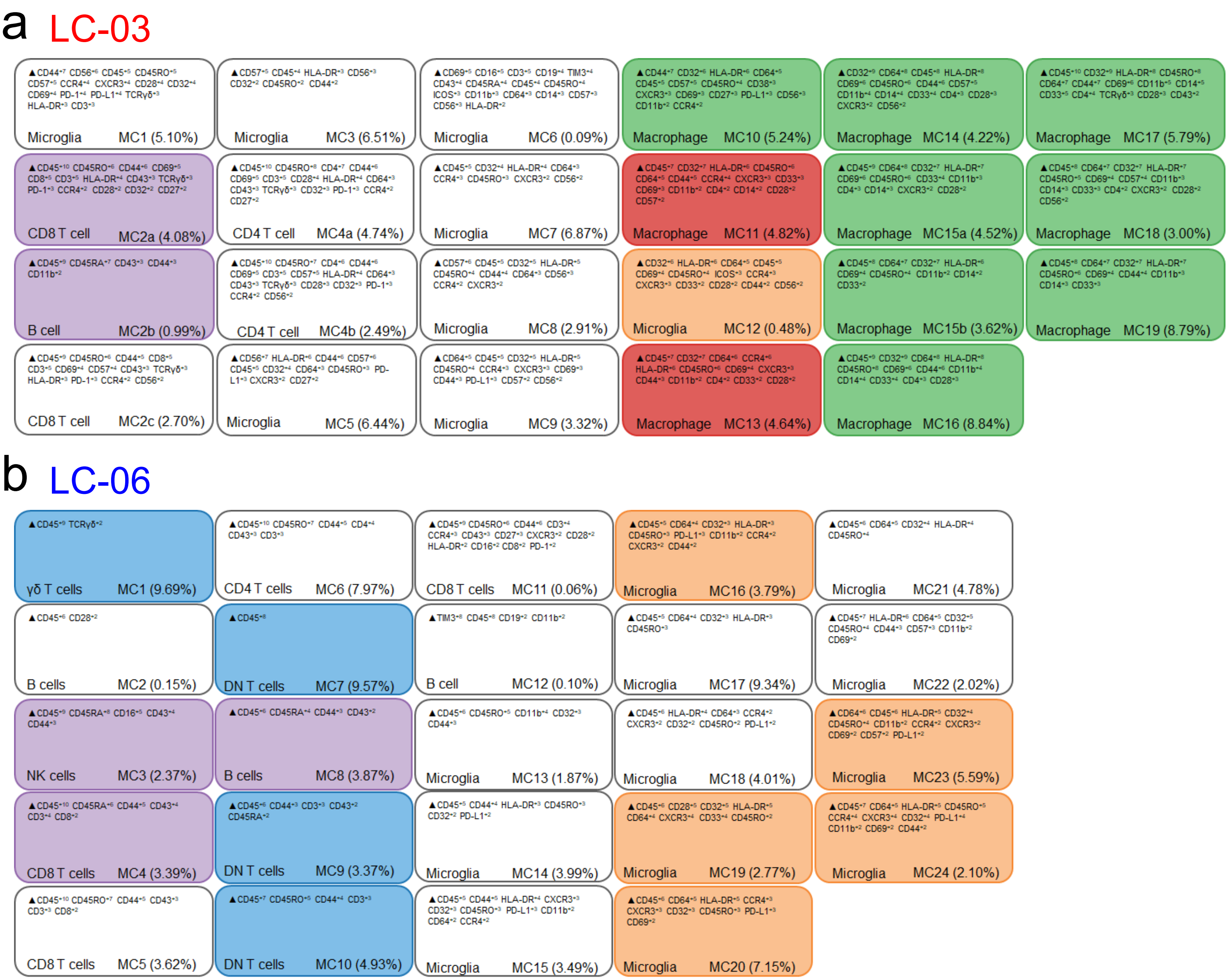


**Supplementary Figure 4: Individual patient MEM labels for each FlowSOM Cluster.**
**a)** MEM labels were generated for each of 19 FlowSOM clusters identified in patient LC-03. **b)** MEM labels generated for each of 24 FlowSOM clusters identified in patient LC-06. Each label is color coded to its corresponding Citrus identity. Percentages in parentheses indicate the frequency of each cluster within the CD45+ fraction in each patient.

Supplementary Figure 5

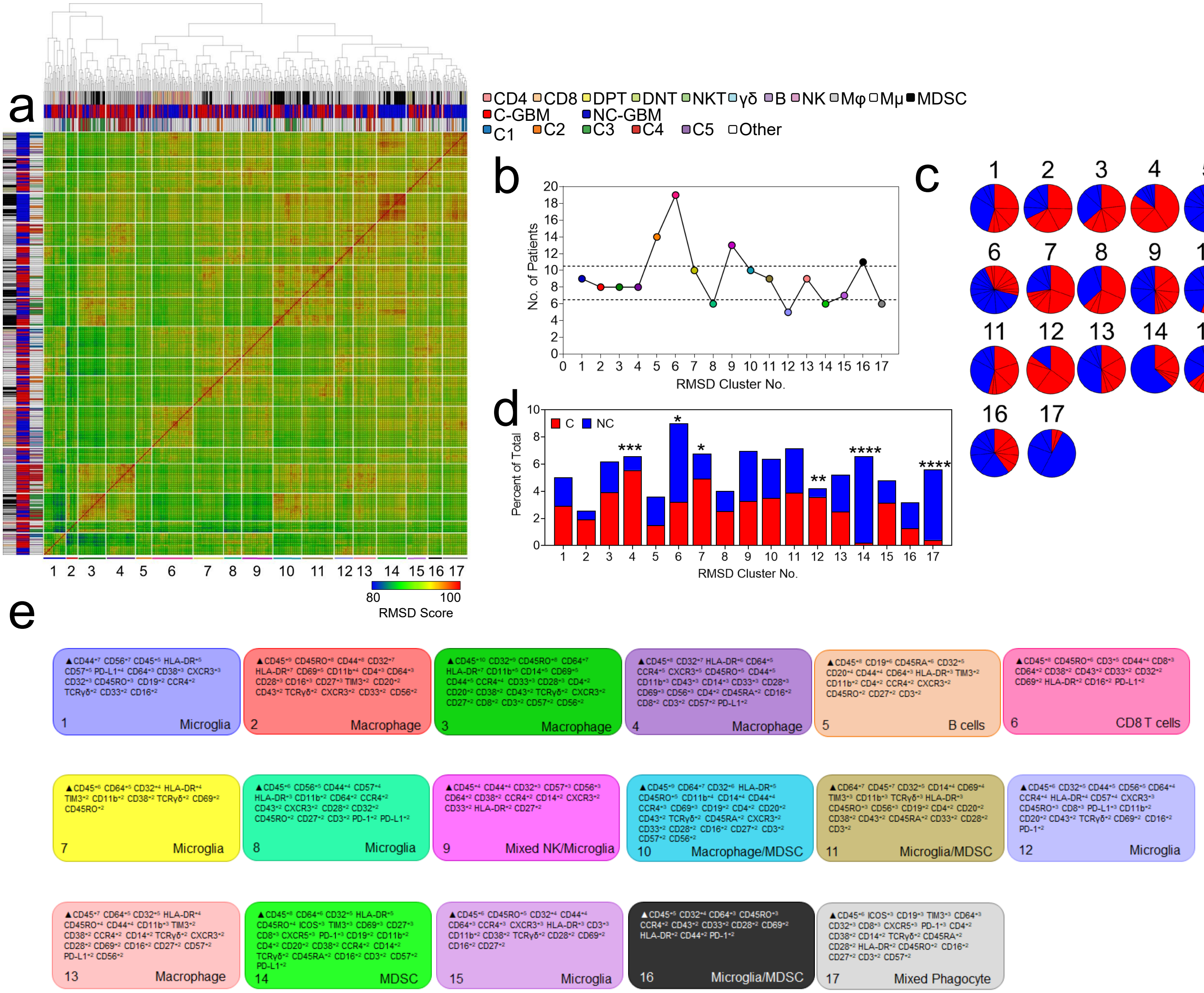


**Supplementary Figure 5: RMSD registration of FlowSOM clusters.**
**a)** Root mean squared deviation (RMSD) was performed to compare the phenotypic similarities between 455 FlowSOM clusters across 19/25 GBM patients, identifying 17 common immune phenotypes. **b)** Graph indicating the number of patients represented within each RMSD metacluster. **c)** Pie charts indicate the proportion of FlowSOM clusters derived from each patient within each indicated RMSD metacluster. **d)** Bar chart indicating the percentage of FlowSOM clusters derived from C-GBM or NC-GBM patients within each RMSD metacluster. **e)** MEM labels from pooled FlowSOM clusters indicate the cellular phenotypes of each larger RMSD metacluster. Red= C-GBM. Blue= NC-GBM. A Chi-Squared analysis determined statistical enrichment. * = $p<0.05$, ** = $p<0.01$, *** = $p<0.001$, **** = $p<0.0001$.

Supplementary Figure 6

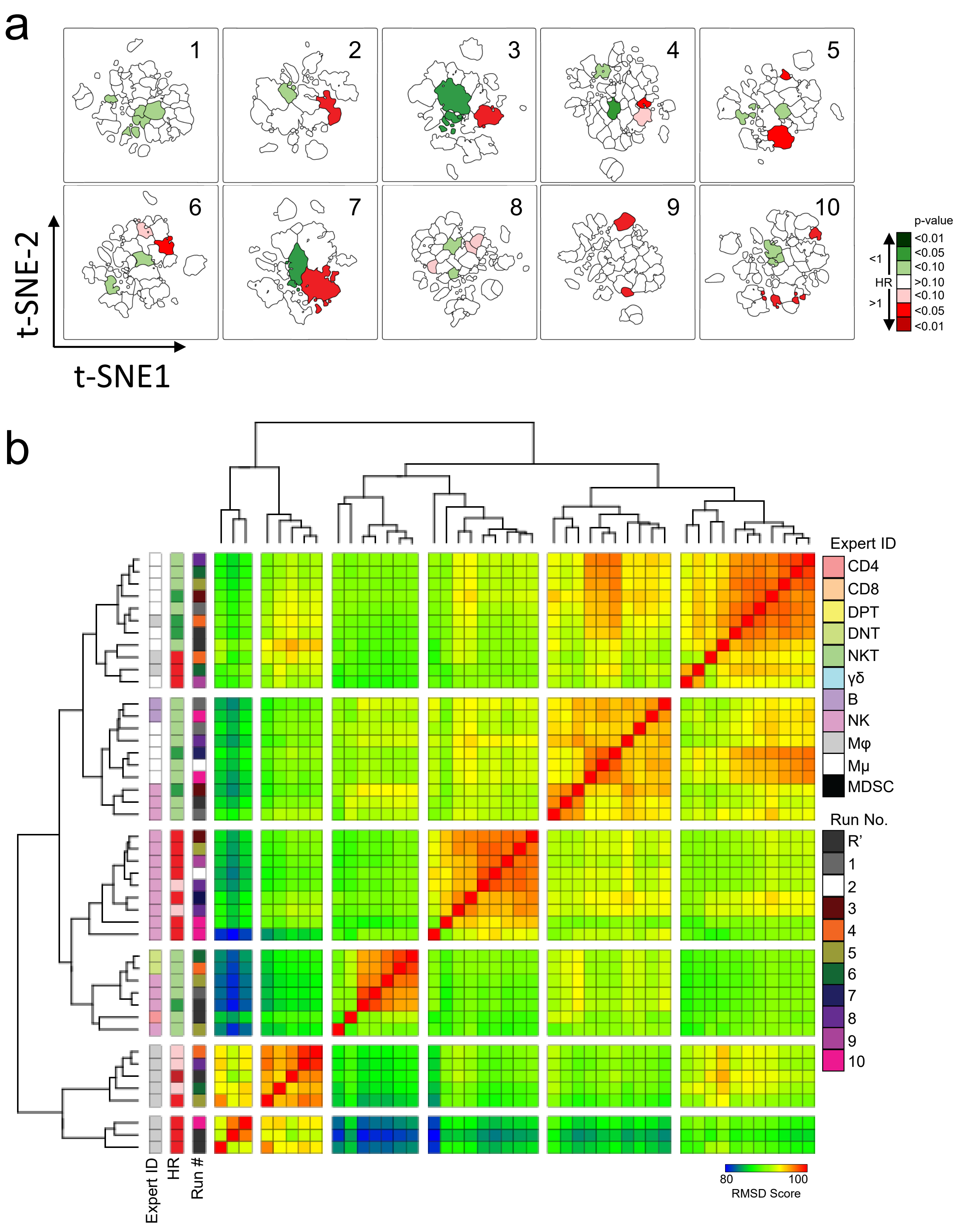


**Supplementary Figure 6: Outcome stratifying immune populations are stably detected by RAPID.** **a)** t-SNE maps indicating FlowSOM clusters whose abundance stratify patient outcome as identified in 10 different implementations of the RAPID algorithm. **b)** RMSD was performed to compare the phenotypic similarities between all risk-stratifying immune populations identified by RAPID. Colors in **a** indicate the hazard associated with more (green) or less (red) favorable outcomes. Colored boxes in **b** indicate the expert identification for each immune subset, hazard ratio, or RAPID iteration respectively.

Supplementary Figure 7

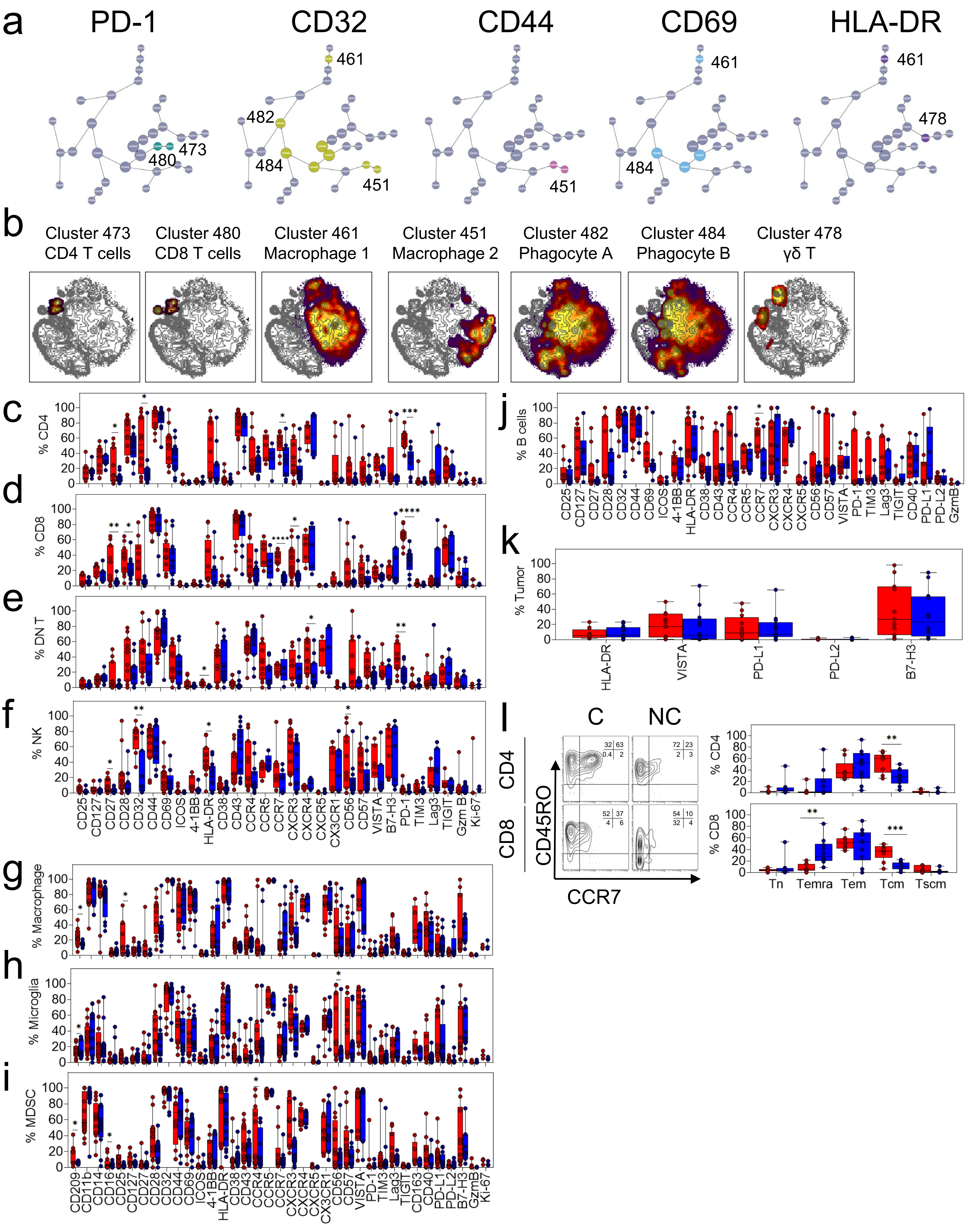

**Supplementary Figure 7: Immune cell phenotypes associated with tumor contact with the lateral ventricle.**

**a)** Citrus clustergrams of immune clusters with statistically different expression of the indicated marker. The most terminal clusters where indicated were chosen for further analysis. **b)** Representative overlay of indicated Citrus clusters (heat) onto the t-SNE map (contour). **c-k)** Box and whisker plots indicating the frequency of CD4 T cells **(c)**, CD8 T cells **(d)**, DNT cells **(e)**, NK cells **(f)**, macrophages **(g)**, microglia **(h)**, myeloid derived suppressor cells **(i)** B cells **(j)** or $CD45^-$ tumor cells **(k)** expressing each indicated marker. **l)** Representative biaxial plots defining the frequency of memory T cells subsets in C- and NC-GBM. Box and whisker plots quantifying the frequency of each memory T cell subset. * = $p<0.05$, ** = $p<0.01$, *** = $p<0.001$, **** = $p<0.0001$.

Supplementary Figure 8

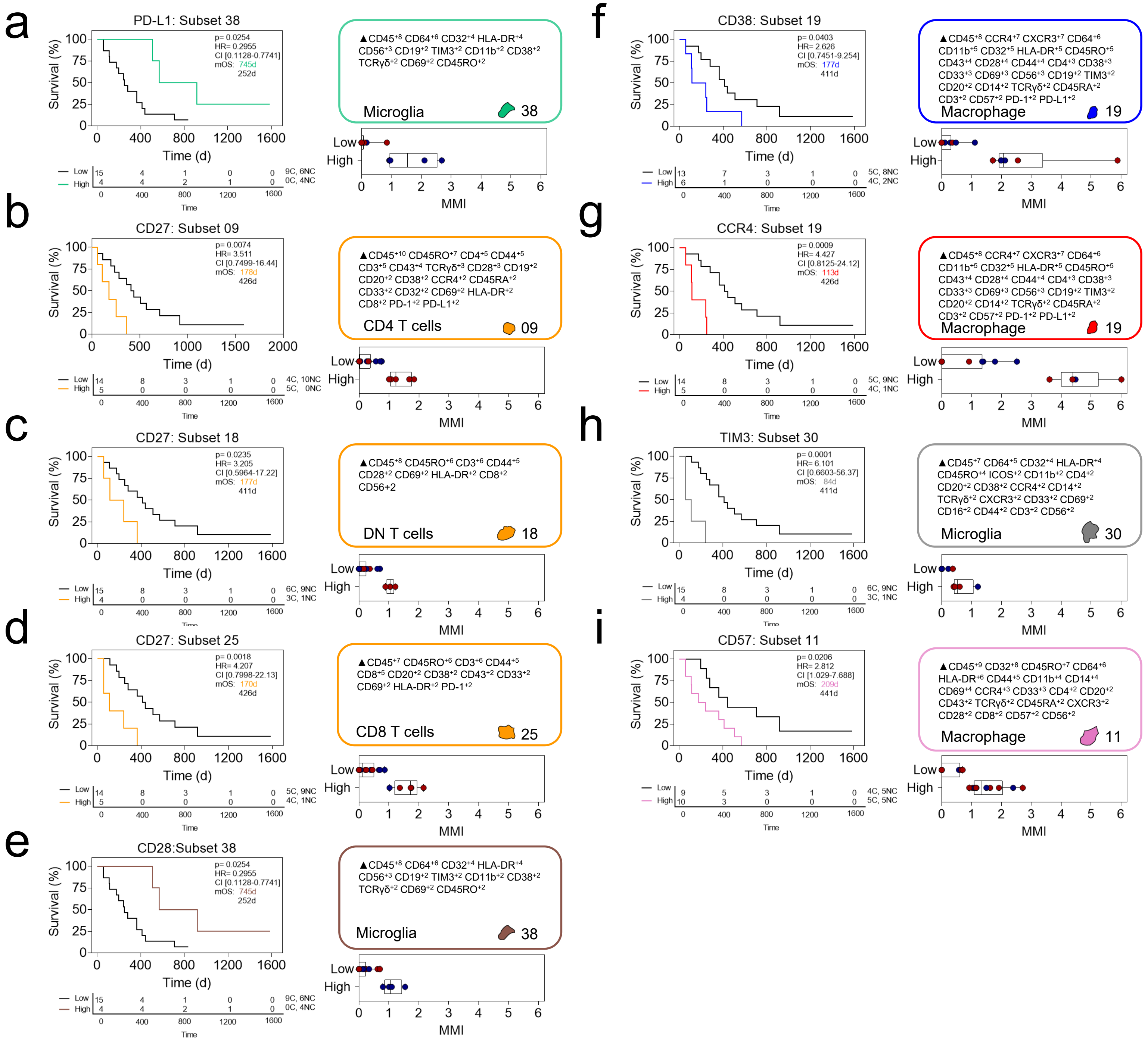


**Supplementary Figure 8: Immune receptor expression correlates with patient outcome.** Kaplan-Meier curves indicating overall survival in immune subsets stratifying patient outcome identified by mmRAPID analysis. Calculated MEM labels identified key features of stratifying immune subsets. Box and whisker diagrams plot the arcsinh transformed median mass intensity for PD-L1 expression **(a)** CD27 **(b-d)**, CD28 **(e)**, CD38 **(f)**, CCR4 **(g)**, TIM3 **(h)**, and CD57 **(i)** in stratified high and low groups.

### Supplementary Figure 9

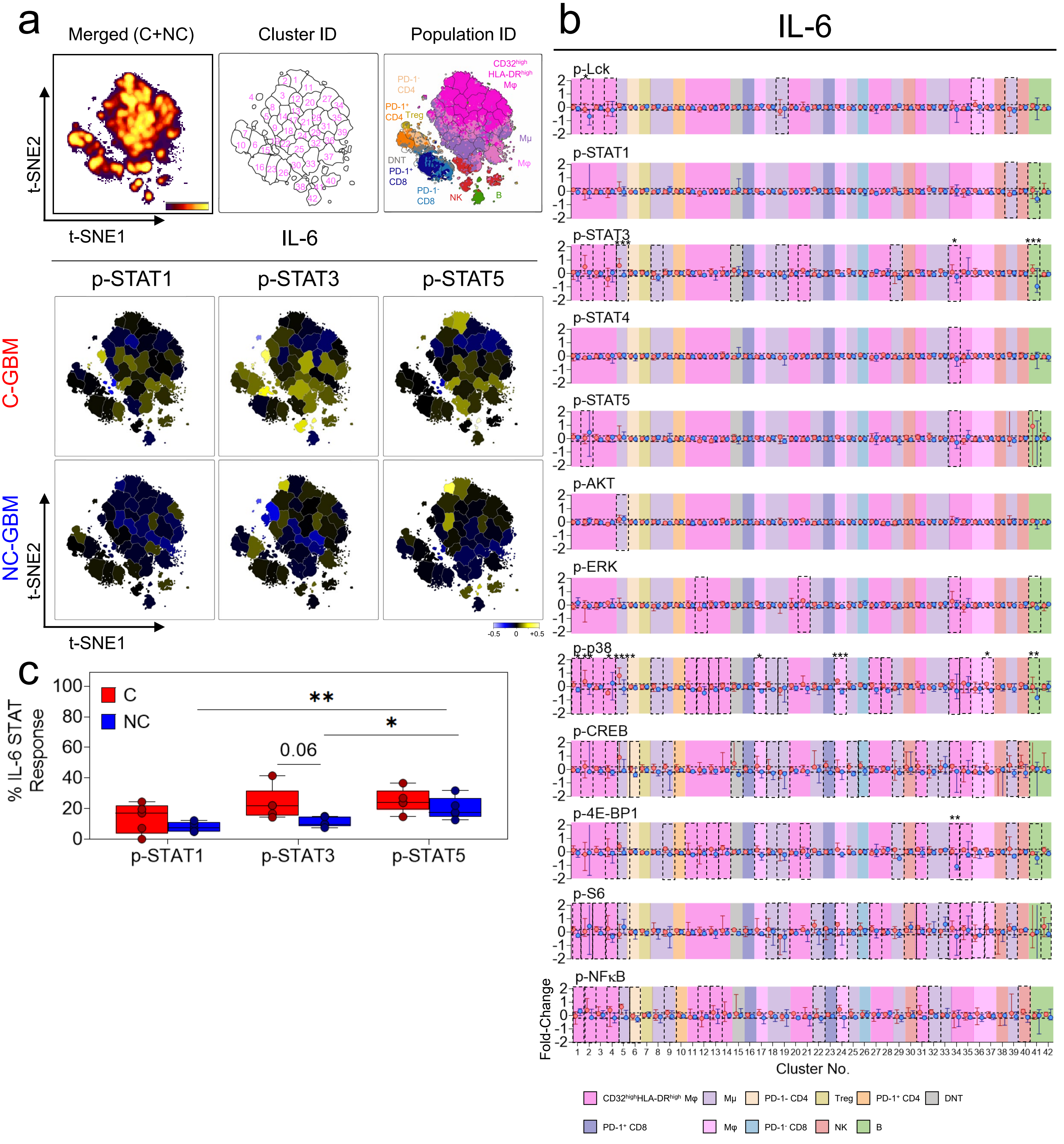

**Supplementary Figure 9: Interleukin 6 induces STAT3 and STAT5 signaling in GBM immune infiltrates.**

**a)** Representative t-SNE plot indicated the density of CD45+ leukocytes (left), enumerated FlowSOM clusters (middle) and overlay of expertly gated immune populations onto the clustered t-SNE axes (right). Representative heatmaps on the t-SNE axes indicate the cluster-specific median arcsinh-fold change of the indicated phosphoprotein under IL-6 stimulation conditions compared to basal phosphorylation. **b)** Graphs indicating the arcsinh transformed phosphoprotein median intensities for each phosphoprotein in each of 42 IL-6 stimulated immune clusters. **c)** Box and whisker plots indicating the proportion of clusters in C-GBM or NC-GBM immune infiltrates surpassing the phospho-signaling threshold (>0.2 arcsinh fold change) in response to IL-6 stimulation. Graphs indicate the median ± IQR. Black horizontal lines in b indicate signaling threshold for each arcsinh transformed value. Black dotted boxes in **b** indicate clusters for which the arcsinh transformed median mass intensity surpassed the signaling threshold (±0.2-fold change) in either the C-GBM or NC-GBM cohort). * = $p<0.05$, ** = $p<0.01$, *** = $p<0.001$, **** = $p<0.0001$.

### Supplementary Figure 10

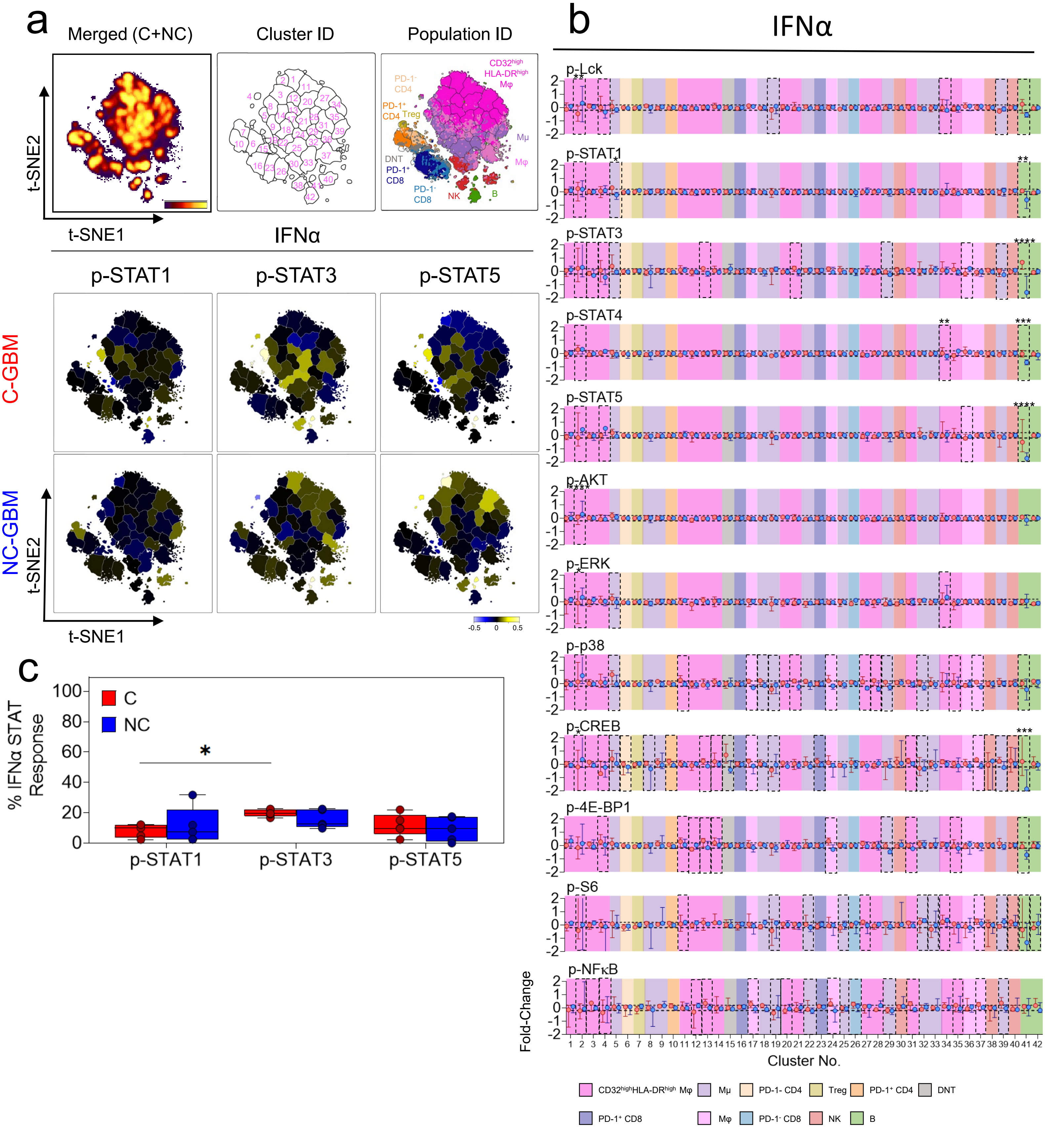

**Supplementary Figure 10: Interferon alpha preferentially induces STAT3 phosphorylation in C-GBM immune infiltrates.**

**a)** Representative t-SNE plot indicated the density of CD45+ leukocytes (left), enumerated FlowSOM clusters (middle) and overlay of expertly gated immune populations onto the clustered t-SNE axes (right). Representative heatmaps on the t-SNE axes indicate the cluster-specific median arcsinh-fold change of the indicated phosphoprotein under IFNα stimulation conditions compared to basal phosphorylation. **b)** Graphs indicating the arcsinh transformed phosphoprotein median intensities for each phosphoprotein in each of 42 IFNα stimulated immune clusters. **c)** Box and whisker plots indicating the proportion of clusters in C-GBM or NC-GBM immune infiltrates surpassing the phospho-signaling threshold (>0.2 arcsinh fold change) in response to IFNα stimulation. Graphs indicate the median ± IQR. Black horizontal lines in b indicate signaling threshold for each arcsinh transformed value. Black dotted boxes in b indicate clusters for which the arcsinh transformed median mass intensity surpassed the signaling threshold (±0.2-fold change) in either the C-GBM or NC-GBM cohort). * = $p<0.05$, ** = $p<0.01$, *** = $p<0.001$, **** = $p<0.0001$.
