## Supplementary Table 1 for "Immune cell regulation in stem cell niche contacting glioblastomas"

**Supplementary Table 1: Patient Demographics and Clinical Characteristics**

**Glioblastoma Patients**

| Sample ID | Gender | Age | Ventricle Contact | Steroids | TMZ | RT | Extent of Resection (subtotal vs. gross total) | IDH Mutation | MGMT promoter methylation | OS (days) | Status | Phenotyping Panel | Checkpoint Panel | Phospho Panel | Citrus Analysis |
|---|---|---|---|---|---|---|---|---|---|---|---|---|---|---|---|
| K01 | F | 59 | Y | Y | Y | Y | STR | WT | N | 364 | dead | X | X | | X |
| LC03 | M | 60 | Y | Y | N | N | STR | WT | N | 57 | dead | X | | | X |
| LC04 | M | 65 | N | Y | Y | Y | GTR | WT | N | 918 | dead | X | X | | X |
| LC06 | M | 41 | N | Y | Y | Y | GTR | WT | N | 1588 | alive | X | X | | X |
| LC08 | F | 55 | N | Y | Y | Y | STR | WT | N | 441 | dead | X | X | X | X |
| LC09 | M | 68 | Y | Y | Y | Y | STR | WT | Y | 110 | dead | X | X | | X |
| LC10 | M | 78 | Y | Y | N | Y | STR | WT | N | 178 | dead | X | X | | X |
| LC11 | F | 76 | N | N | Y | Y | STR | WT | Y | 411 | dead | X | X | | X |
| LC12 | M | 40 | N | Y | Y | Y | GTR | WT | Y | 710 | dead | X | X | | X |
| LC18 | M | 69 | N | Y | Y | Y | STR | WT | N | 240 | dead | X | X | | X |
| LC22 | M | 64 | Y | Y | Y | Y | STR | WT | N | 366 | dead | X | X | X | X |
| LC25 | F | 60 | N | N | Y | Y | GTR | WT | N | 731 | dead | X | X | | |
| LC26 | M | 71 | N | Y | Y | Y | STR | WT | Y | 836 | alive | X | X | | X |
| LC27 | M | 62 | Y | Y | Y | Y | STR | WT | N | 252 | dead | X | X | X | X |
| LC36 | F | 66 | N | Y | Y | Y | STR | WT | N | 317 | dead | X | X | | |
| RT01 | F | 69 | Y | Y | Y | Y | STR | WT | N | 198 | dead | X | X | X | X |
| RT07 | F | 70 | Y | Y | Y | Y | STR | WT | N | 57 | dead | X | X | | X |
| RT10 | M | 56 | Y | Y | N | N | STR | WT | Y | 113 | dead | X | X | X | X |
| RT14 | F | 55 | N | Y | Y | Y | GTR | WT | Y | 571 | dead | X | X | X | X |
| RT15 | F | 80 | Y | Y | N | Y | STR | WT | N | 353 | dead | X | X | X | |
| W02 | M | 75 | N | Y | Y | Y | STR | WT | Y | 507 | dead | X | X | X | X |
| W05 | M | 60 | N | Y | Y | Y | STR | WT | N | 282 | dead | X | X | X | X |
| W11 | M | 69 | N | Y | Y | Y | GTR | WT | N | 896 | dead | X | X | X | |
| W12 | M | 50 | Y | Y | Y | Y | STR | WT | Y | 723 | dead | X | X | | |
| W14 | M | 47 | Y | Y | Y | Y | STR | WT | Y | 488 | dead | X | X | | |

**Healthy Donor PBMC**

| | | | | | | | | | | | | | | |
|---|---|---|---|---|---|---|---|---|---|---|---|---|---|---|
| VU026 | | | | | | | | | | | | X | | |
| VU027 | | | | | | | | | | | | X | | |
| VU043 | | | | | | | | | | | | | X | X |
