## Supplementary Table 2 for "Immune cell regulation in stem cell niche contacting glioblastomas"

**Supplementary Table 2: Mass Cytometry Immune Phenotyping Panel**

| Metal | Mass | Marker | Clone | Vendor | Catalog Number | Citrus Abundance Clustering | Citrus Median Clustering | Citrus Median Expression |
|---|---|---|---|---|---|---|---|---|
| Y | 89 | | | | | | | |
| Pd | 102 | | | | | | | |
| Rh | 103 | Live/Dead | | Fluidigm | 201103A | | | |
| Pd | 104 | | | | | | | |
| Pd | 105 | | | | | | | |
| Pd/Cd | 106 | | | | | | | |
| Cd | 108 | | | | | | | |
| Pd/Cd | 110 | | | | | | | |
| Cd | 111 | | | | | | | |
| Cd | 112 | | | | | | | |
| Cd | 113 | | | | | | | |
| Cd | 114 | | | | | | | |
| Cd | 116 | | | | | | | |
| I | 127 | | | | | | | |
| La | 139 | | | | | | | |
| Pr | 141 | ICOS | C398.4A | Biolegend | 313502 | X | | X |
| Nd | 142 | CD19 | HIBI-9 | Fluidigm | 3142001B | X | X | |
| Nd | 143 | TIM3 | F38-232 | Biolegend | 345019 | X | | X |
| Nd | 144 | CD11b | ICRF44 | Fluidigm | 3144001B | X | X | |
| Nd | 145 | CD4 | RPA-T4 | Fluidigm | 3145001B | X | X | |
| Nd | 146 | CD64 | 10.1 | Fluidigm | 3146006B | X | X | |
| Sm | 147 | CD20 | 2H7 | Fluidigm | 3147001B | X | X | |
| Nd | 148 | CD38 | HIT2 | Biolegend | 303535 | X | | X |
| Sm | 149 | CCR4 | L291H4 | Fluidigm | 3149029A | X | | X |
| Nd | 150 | CD43 | 84-3C1 | Fluidigm | 3150006B | X | | X |
| Eu | 151 | CD14 | M5E2 | Fluidigm | 3151009B | X | X | X |
| Sm | 152 | TCRgd | 11F2 | Fluidigm | 3152008B | X | X | |
| Eu | 153 | CD45RA | HI100 | Fluidigm | 3152001B | X | X | |
| Sm | 154 | CD45 | HI30 | Fluidigm | 3154001C | X | X | |
| Gd | 155 | | | | | | | |
| Gd | 156 | CXCR3 | G025H7 | Fluidigm | 3156004B | X | | X |
| Gd | 158 | CD33 | WM53 | Fluidigm | 3158001B | X | X | |
| Tb | 159 | CCR7 | G043H7 | Fluidigm | 3159003A | X | | X |
| Gd | 160 | CD28 | CD28.2 | Fluidigm | 3160003B | X | | X |
| Dy | 161 | CD32 | FUN2 | Biolegend | 303202 | X | | X |
| Dy | 161 | Ki-67 | B56 | Fluidigm | 3160002B | | | |
| Dy | 162 | CD69 | FN50 | Fluidigm | 3162001C | X | | X |
| Dy | 163 | HLA-DR | L243 | Biolegend | 307651 | X | | X |
| Dy | 164 | CD45RO | UCHL1 | Fluidigm | 3164007B | X | X | |
| Ho | 165 | CD16 | 3G8 | Fluidigm | 3165001B | X | X | |
| Er | 166 | CD44 | BJ18 | Fluidigm | 3166001C | X | | X |
| Er | 167 | CD27 | O323 | Fluidigm | 3167002B | X | | X |
| Er | 168 | CD8 | SK1 | Fluidigm | 3168002B | X | X | |
| Tm | 169 | CD25 | 2A3 | Fluidigm | 3169003B | X | | |
| Er | 170 | CD3 | UCHT1 | Fluidigm | 317001C | X | X | |
| Yb | 171 | CXCR5 | RF8B2 | Fluidigm | 3171014B | X | X | |
| Yb | 172 | CD57 | HCD57 | Fluidigm | 3172009B | X | | X |
| Yb | 173 | Granzyme B | GB11 | Fluidigm | 3171002B | | | |
| Yb | 174 | PD-1 | EH12.2H7 | Fluidigm | 3174020B | X | | X |
| Lu | 175 | PD-L1 | 28E.2A3 | Fluidigm | 3175017B | X | | X |
| Yb | 176 | CD56 | CMSSB | Fluidigm | 3176003B | X | X | |
| Ir | 191 | Intercalator | | Fluidigm | 201192B | | | |
| Ir | 193 | | | | | | | |
| Pt | 194 | | | | | | | |
| Pt | 195 | | | | | | | |
| Pt | 196 | | | | | | | |
| Pt | 198 | | | | | | | |
| Bi | 209 | | | | | | | |
