## Supplementary Table 3 for "Immune cell regulation in stem cell niche contacting glioblastomas"

| Supplementary Table 3: Mass Cytometry Immune Checkpoint Receptor Panel | | | | | |
|---|---|---|---|---|---|
| **Metal** | **Mass** | **Marker** | **Clone** | **Vendor** | **Catalogue Number** |
| Y | 89 | CD45 | HI30 | Fluidigm | 3089003B |
| Pd | 102 | | | | |
| Rh | 103 | Live/Dead | | Fluidigm | 201103A |
| Pd | 104 | | | | |
| Pd | 105 | | | | |
| Pd/Cd | 106 | | | | |
| Cd | 108 | | | | |
| Pd/Cd | 110 | | | | |
| Cd | 111 | | | | |
| Cd | 112 | | | | |
| Cd | 113 | | | | |
| Cd | 114 | | | | |
| Cd | 116 | | | | |
| I | 127 | | | | |
| La | 139 | | | | |
| Pr | 141 | CD209 | 14E3G7 | R&D Systems | 3999999-2 |
| Nd | 142 | CD19 | HIBI-9 | Fluidigm | 3142001B |
| Nd | 143 | CD127 | A019D55 | Fluidigm | 3143012B |
| Nd | 144 | CD11b | ICRF44 | Fluidigm | 3144001B |
| Nd | 145 | CD4 | RPA-T4 | Fluidigm | 3145001B |
| Nd | 146 | CD64 | 10.1 | Fluidigm | 3146006B |
| Sm | 147 | CD20 | 2H7 | Fluidigm | 3147001B |
| Nd | 148 | CD38 | HIT2 | Biolegend | 303535 |
| Sm | 149 | CCR4 | L291H4 | Fluidigm | 3149029A |
| Nd | 150 | Lag3 | 874501 | Fluidigm | 3150016B |
| Eu | 151 | CD14 | M5E2 | Fluidigm | 3151009B |
| Sm | 152 | | | | |
| Eu | 153 | TIGIT | MBSA43 | Fluidigm | 3153019B |
| Sm | 154 | PD-L2 | 24F.10C12 | Biolegend | 329602 |
| Gd | 155 | RGMb | 298528 | R&D Systems | MAB3630 |
| Gd | 156 | CXCR3 | G025H7 | Fluidigm | 3156004B |
| Gd | 158 | 4-1BB | 4B4-1 | Fluidigm | 3158013B |
| Tb | 159 | CCR7 | G043H7 | Fluidigm | 3159003A |
| Gd | 160 | VISTA | D1L2G | Fluidigm | 3160025D |
| Dy | 161 | CD32 | FUN2 | Biolegend | 303202 |
| Dy | 162 | CD69 | FN50 | Fluidigm | 3162001C |
| Dy | 163 | HLA-DR | L243 | Biolegend | 307651 |
| Dy | 164 | CD45RO | UCHL1 | Fluidigm | 3164007B |
| Ho | 165 | CD40 | 5C3 | Fluidigm | 3165005B |
| Er | 166 | CD44 | BJ18 | Fluidigm | 3166001C |
| Er | 167 | CD27 | O323 | Fluidigm | 3167002B |
| Er | 168 | CD8 | SK1 | Fluidigm | 3168002B |
| Tm | 169 | CD25 | 2A3 | Fluidigm | 3169003B |
| Er | 170 | CD3 | UCHT1 | Fluidigm | 317001C |
| Yb | 171 | Granzyme B | GB11 | Fluidigm | 3171002B |
| Yb | 172 | CX3CR1 | 2A9-1 | Fluidigm | 3172017B |
| Yb | 173 | B7-H3 | polyclonal | Fluidigm | 3173014D |
| Yb | 174 | PD-1 | EH12.2H7 | Fluidigm | 3174020B |
| Lu | 175 | PD-L1 | 28E.2A3 | Fluidigm | 3175017B |
| Yb | 176 | CD56 | CMSSB | Fluidigm | 3176003B |
| Ir | 191 | Intercalator | | Fluidigm | 201192B |
| Ir | 193 | | | | |
| Pt | 194 | | | | |
| Pt | 195 | | | | |
| Pt | 196 | | | | |
| Pt | 198 | | | | |
| Bi | 209 | CD16 | 3G8 | Fluidigm | 3209002B |
