## Supplementary Table 4 for "Immune cell regulation in stem cell niche contacting glioblastomas"

| Supplementary Table 4: Mass Cytometry Immune Phospho-signaling Panel | | | | | |
|---|---|---|---|---|---|
| **Metal** | **Mass** | **Marker** | **Clone** | **Vendor** | **Catalogue Number** |
| Y | 89 | CD45 | HI30 | Fluidigm | 3089003B |
| Pd | 102 | | | | |
| Rh | 103 | Live/Dead | | Fluidigm | 201103A |
| Pd | 104 | | | | |
| Pd | 105 | | | | |
| Pd/Cd | 106 | | | | |
| Cd | 108 | | | | |
| Pd/Cd | 110 | | | | |
| Cd | 111 | | | | |
| Cd | 112 | | | | |
| Cd | 113 | | | | |
| Cd | 114 | | | | |
| Cd | 116 | | | | |
| I | 127 | | | | |
| La | 139 | | | | |
| Pr | 141 | CD11b | ICRF44 | Biolegend | 301337 |
| Nd | 142 | CD19 | HIBI-9 | Fluidigm | 3142001B |
| Nd | 143 | CD127 | A019D55 | Fluidigm | 3143012B |
| Nd | 144 | CCR5 | NP-6G4 | Fluidigm | 3144007A |
| Nd | 145 | CD4 | RPA-T4 | Fluidigm | 3145001B |
| Nd | 146 | CD64 | 10.1 | Fluidigm | 3146006B |
| Sm | 147 | CD20 | H1 | Fluidigm | 3147001B |
| Nd | 148 | CD38 | HIT2 | Biolegend | 303535 |
| Sm | 149 | p-4EBP1 (T37/46) | 236B4 | Fluidigm | 3149005A |
| Nd | 150 | p-STAT5 (Y694) | 47 | Fluidigm | 3150005A |
| Eu | 151 | CD14 | M5E2 | Fluidigm | 3151009B |
| Sm | 152 | p-AKT (S743) | D9E | Fluidigm | 3152005A |
| Eu | 153 | p-STAT1 (Y701) | 58D6 | Fluidigm | 3153003A |
| Sm | 154 | CD163 | GHI/61 | Fluidigm | 3154007B |
| Gd | 155 | CD45RA | HI100 | Fluidigm | 3155011B |
| Gd | 156 | p-p38 (T180/Y182) | D3F9 | Fluidigm | 3156002A |
| Gd | 158 | p-STAT3 (Y705) | 4/P-STAT3 | Fluidigm | 3158005A |
| Tb | 159 | CCR7 | G043H7 | Fluidigm | 3159003A |
| Gd | 160 | CD28 | CD28.2 | Fluidigm | 3160003B |
| Dy | 161 | CD32 | FUN2 | Biolegend | 303202 |
| Dy | 162 | p-Lck (T505) | 4/LCK-Y505 | Fluidigm | 3162004A |
| Dy | 163 | HLA-DR | L243 | Biolegend | 307651 |
| Dy | 164 | CD45RO | UCHL1 | Fluidigm | 3164007B |
| Ho | 165 | p-CREB | 87G3 | Fluidigm | 3165009A |
| Er | 166 | p-NFκBp65 (S529) | K10-895.12.50 | Fluidigm | 3166006A |
| Er | 167 | CD27 | O323 | Fluidigm | 3167002B |
| Er | 168 | CD8 | SK1 | Fluidigm | 3168002B |
| Tm | 169 | CD25 | 2A3 | Fluidigm | 3169003B |
| Er | 170 | CD3 | UCHT1 | Fluidigm | 317001C |
| Yb | 171 | p-ERK1/2 (T202/Y204) | D13.14.4E | Fluidigm | 3167005A |
| Yb | 172 | p-S6 (S235/236) | N7-548 | Fluidigm | 3172008A |
| Yb | 173 | CXCR4 | 12G5 | Fluidigm | 3173001B |
| Yb | 174 | p-STAT4 (Y693) | 38/p-STAT4 | Fluidigm | 3174005A |
| Lu | 175 | PD-1 | EH12.2H7 | Fluidigm | 3175008B |
| Yb | 176 | CD56 | CMSSB | Fluidigm | 3176003B |
| Ir | 191 | Intercalator | | Fluidigm | 201192B |
| Ir | 193 | | | | |
| Pt | 194 | | | | |
| Pt | 195 | | | | |
| Pt | 196 | | | | |
| Pt | 198 | | | | |
| Bi | 209 | CD16 | 3G8 | Fluidigm | 3209002B |
